# Large-scale phylogenomics of aquatic bacteria reveal molecular mechanisms for adaptation to salinity

**DOI:** 10.1101/2022.10.03.510577

**Authors:** Krzysztof T Jurdzinski, Maliheh Mehrshad, Luis Fernando Delgado, Ziling Deng, Stefan Bertilsson, Anders F Andersson

**Author notes:** Equal contribution (Krzysztof T Jurdzinski, Maliheh Mehrshad).

## Abstract

The crossing of environmental barriers poses major adaptive challenges. Rareness of freshwater-marine transitions separates the bacterial communities, but how these are related to brackish counterparts remains elusive, as are molecular adaptations facilitating cross-biome transitions. Here, we conduct large-scale phylogenomic analysis of freshwater, brackish, and marine quality-filtered metagenome-assembled genomes (11,276 MAGs). Average nucleotide identity analyses showed that bacterial species rarely existed in multiple biomes. Distinct brackish basins co-hosted numerous species despite differences in salinity and geographic distance, the latter having stronger intra-species population structuring effects. We further identified the most recent cross-biome transitions, which were rare, ancient, and most commonly directed towards the brackish biome. Transitions were accompanied by changes in isoelectric point distribution and amino acid composition of inferred proteomes, as well as convergent gains or losses of specific gene functions. Therefore, adaptive challenges entailing proteome reorganization and specific changes in gene content result in species-level separation between aquatic biomes.

## Introduction

Transitions between different environmental conditions have long been identified as one of the major drivers of evolutionary innovation^1^. Evidence of fast dispersal of bacteria across the globe suggests that physicochemical and biological factors constrain the distribution of bacteria to a higher extent than geography and dispersal limitation^2^. However, the highly dynamic nature of bacterial genomes^3^, including the ability to acquire new genes through horizontal gene transfer^4^, can potentially facilitate the crossing of environmental barriers.

Salinity, pH, temperature, and other factors important for structuring microbial diversity are predicted to change in coastal waters due to climate change^5^. Combined with the crucial role of bacteria in such ecosystems^6^, there is a need to understand the potential and limitations of adaptation as well as dispersal in response to changing environmental conditions. Comparative genome analyses across major environmental barriers have the potential to uncover the molecular mechanisms of such adaptation, as phylogenetic frameworks allow to disentangle the effects of habitat from differences in taxonomic composition.

Salinity has long been considered the strongest physicochemical factor determining bacterial community composition^7,8^. Distinct bacterial lineages inhabit freshwater and marine biomes, and transitions between these biomes have been rare in the documented evolutionary history of aquatic bacteria^9–11^, pinpointing differences in salinity as a particularly strong barrier for microbes to overcome. More recently, fragment-recruitment-based metagenomic analyses have indicated that brackish environments with intermediate salinity levels also host genetically distinct bacterial lineages^12,13^. However, this has not been confirmed in a rigorous phylogenomic framework and little is known about the biogeography of the potential brackish microbiome and its relationship with its freshwater and marine counterparts. Many questions remain unanswered, such as which groups of bacteria have transitioned between the aquatic biomes, how often, in which directions, and when have the transitions taken place.

Protein properties such as isoelectric points were recently shown to differ between aquatic biomes with a pattern that may reflect adaptation to different extra- and intracellular ion concentrations^14^. Detected changes are mainly seen for surfaces of soluble proteins and differ between certain groups of closely related freshwater, brackish, and marine lineages^14^. Likewise, the functional gene content of microbial communities changes in a salinity-dependent manner^11,15–17^, though it is unclear to what extent these effects are merely attributable to differences in the taxonomic composition. The widespread patterns of convergent gene gain and loss accompanying the cross-biome transition, as well as evolutionary factors and dynamics driving the shift in isoelectric point distribution, could be further elucidated through a systematic phylogenomic analysis across the bacterial tree of life.

Here we analyze an extensive and diverse set of freshwater, brackish, and marine metagenome-assembled genomes (MAGs) in a phylogenomic framework. We cluster MAGs by average nucleotide identity (ANI) and compare the similarities of genomes from the same and different biomes. We identify the most recent transitions between the aquatic biomes and estimate their evolutionary timeline and directionality. Finally, we show that large-scale changes in the properties of the predicted proteome and specific alterations in gene content accompany the transitions in diverse bacterial lineages.

## Results

### A comprehensive dataset of MAGs from freshwater, brackish and marine biomes

A total of 13,783 MAGs were collected from freshwater^18–20^, brackish^17,21^, and marine^22,23^ biomes (Fig 1a) to analyze evolutionary histories and functional adaptations in aquatic bacteria. The MAGs were filtered at ≥75% completeness and ≤5% contamination thresholds, and the 11,509 MAGs passing these criteria were considered for the remaining analyses (11,276 bacterial and 233 archaeal). For further analyses, we focused on the bacterial MAGs, of which 7643 were from freshwater, 2268 from brackish, and 1365 from marine waters (Table 1). Collected MAGs contained representatives from 72 phyla distributed across 135 classes and 348 orders, with the majority (91%) belonging to only 12 bacterial phyla with >100 representatives per phylum (Supplementary Data S1).

**Fig. 1.**
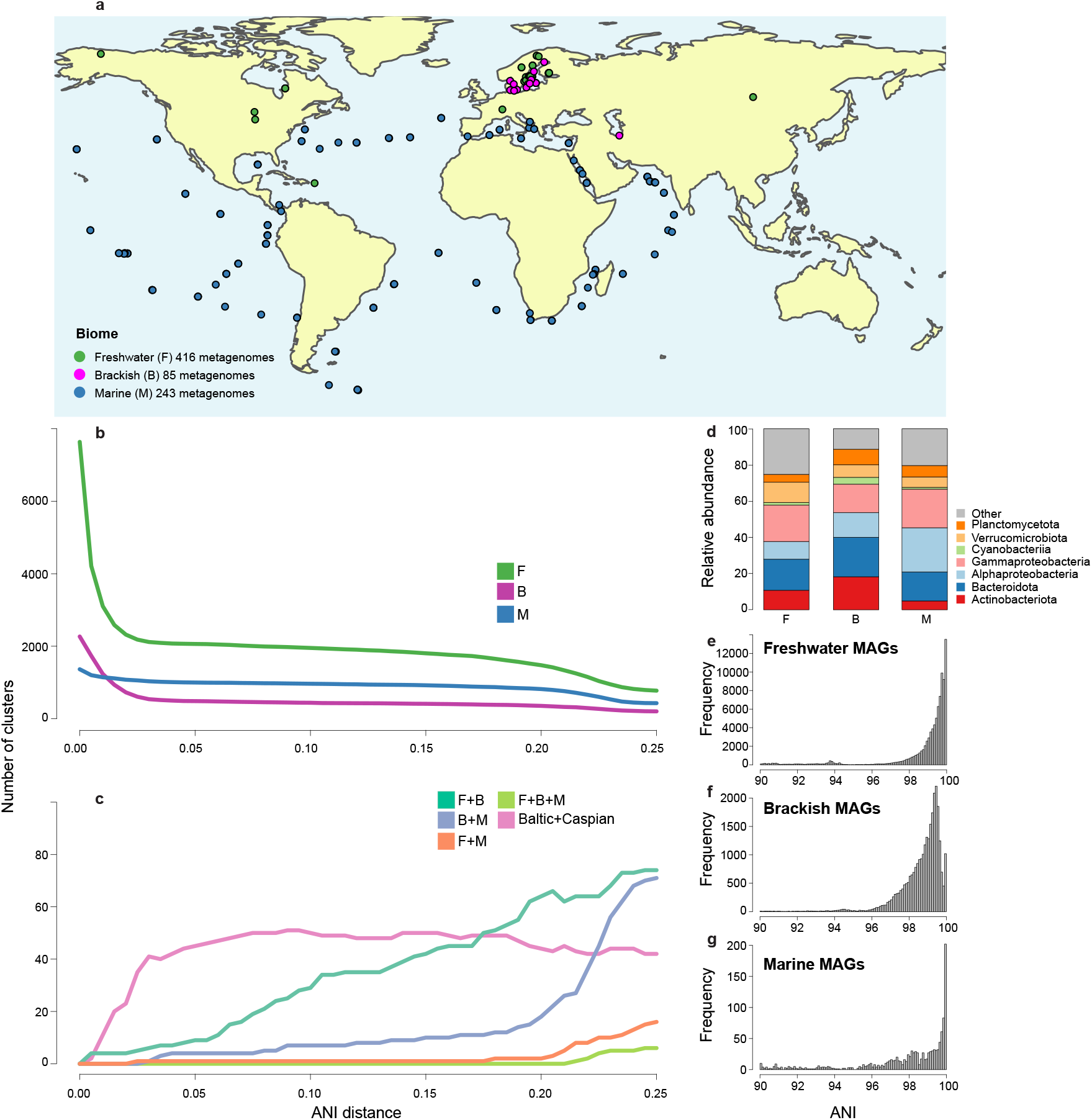
Status of shared bacterial clusters in different biomes. a. Map showing origins of MAGs [refs to papers from where we got the MAGs] **b.** Number of MAG clusters at different nucleotide distance cut-offs in each biome. **c.** Number of MAG clusters at different nucleotide distance cut-offs with members of different biome combinations. **d**. relative abundance of MAGs affiliated to different phyla in the three different biomes. Frequency distributions of pairwise inter-MAG ANI values in the range of 90 – 100% in **e.** freshwater, **f.** brackish, and **g.** marine genomes.

**Table 1.** Number of MAGs originated from each dataset

| Dataset | #metagenomes | #Total MAGs | #filtered MAGs | #filtered Bacteria | #filtered Archaea | Reference |
| --- | --- | --- | --- | --- | --- | --- |
| North American lakes (F) | 141 | 193 | 84 | 84 | 0 | Linz et al. (2018) |
| StratfreshDB (F) | 267 | 8554 | 7654 | 7554 | 100 | Buck et al. (2021) |
| Lake Baikal (F) | 2 | 35 | 7 | 5 | 2 | Cabello-Yeves et.al. (2018) |
| Baltic Sea (B) | 82 | 1989 | 1989 | 1954 | 35 | Alneberg et al. (2020) |
| Caspian Sea (B) | 3 | 324 | 324 | 314 | 10 | Mehrshad et al. (2016) |
| Global Ocean T (M) | 243* | 2631 | 990 | 932 | 58 | Tully et al. (2018) |
| Global Ocean D (M) | 243* | 957 | 461 | 433 | 28 | Delmont et al. (2017) |

\* The same metagenomes were used to obtain MAGs by Delmont et al. (2017) and Tully et al. (2018) using different approaches.

### Bacterial species are rarely shared between biomes

Clustering the bacterial MAGs using ANI at the threshold of the operational definition of species (95%^24,25^) resulted in 3561 genome clusters (Fig 1b). MAGs of the vast majority of genome clusters (n=3547) belonged to a single biome (2063 freshwater, 485 brackish, and 999 marine genome clusters), while no genome cluster had representatives from all three biomes. Only fourteen genome clusters harbored MAGs belonging to pairs of biomes (nine, four, and one genome clusters with freshwater-brackish (FB), brackish-marine (BM), and freshwater-marine (FM) MAGs, respectively) (Fig 1c). The only shared FM genome cluster was affiliated to the genus *Limonbacter* (phylum Proteobacteria). Shared FB clusters were affiliated to phyla Actinobacteriota, Proteobacteria, Verrucomicrobiota, and Bacteroidota. The shared BM clusters belonged to phyla Proteobacteria, Verrucomicrobiota, and Bacteroidota.

Fortyfive of the 485 brackish genome clusters had members from both included brackish water bodies; the Baltic and the Caspian Sea, while 300 and 141 were unique to Baltic and Caspian, respectively. The shared clusters belonged to 13 classes in 10 different phyla (Planctomycetota, Verrucomicrobiota, Gemmatimonadota, Bacteroidota, Actinobacteriota, Chloroflexota, Cyanobacteria, Firmicutes, Nitrospinota, and Proteobacteria). The extent of overlap between Baltic and Caspian Sea genome clusters (18.9%) was significantly higher than between brackish and freshwater (0.007%) and brackish and marine clusters (0.012%; *P* < 10^−16^ for both comparisons), despite the geographic separation and lack of hydrological connectivity between these ecosystems. This supports the previously postulated existence of a global brackish-specific microbiome^12,21^.

The clustering results above indicate that bacterioplankton belonging to the same species (within the 95% ANI threshold) are inhabiting the geographically distinct and distant Baltic and Caspian Seas. To investigate intra-specific (within-species) biogeographical patterns, we used a population genomics approach, where allele frequencies in single nucleotide variant (SNV) loci are obtained by aligning metagenomic sequences to the genomes. As previously reported^26^, principal coordinates (PCoA) analysis of pairwise fixation indices (*F_ST_*) showed that for each of the five genome clusters where we applied this approach in the present study, populations were highly structured according to salinity within the Baltic Sea (Fig. 2a). However, adding the Caspian Sea data to the analysis revealed even stronger population structuring between the two seas (Fig. 2b), indicating geographic separation at the strain level within these species.

**Fig. 2.**
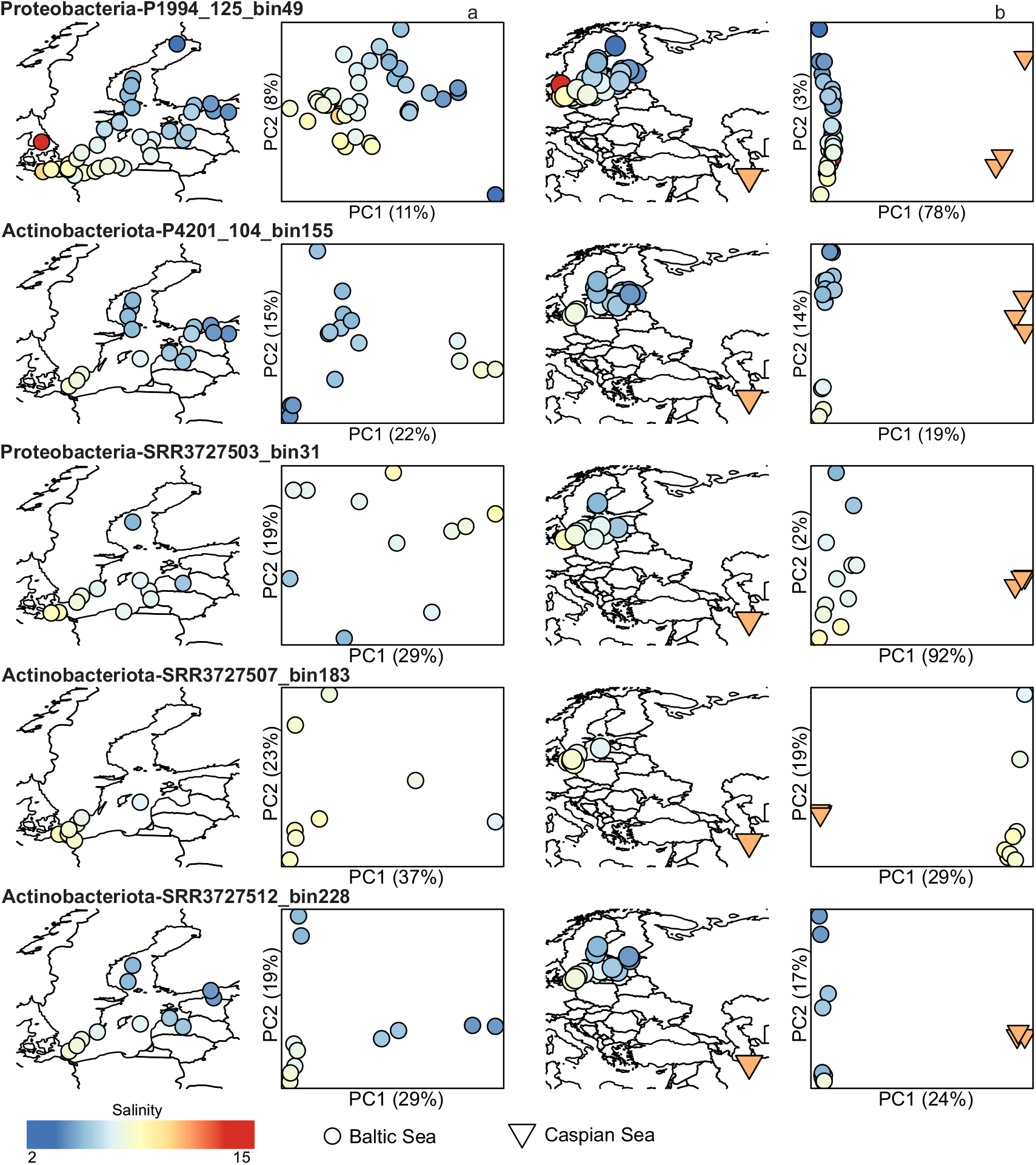
Population structure of brackish genome clusters. a-b, Geographic origin of each metagenome sample included for the MAG is shown to the left and a PCoA based on pairwise *F_ST_* values to the right. Only Baltic Sea samples were included in **a**. whereas both Baltic Sea and Caspian Sea samples were included in **b.**

### Transitions between aquatic biomes are rare, ancient, and in most cases directed into the brackish biome

We reconstructed the phylogeny of the aquatic MAGs using a set of conserved housekeeping genes^27^. The resulting tree was pruned to contain a maximum of one MAG from each biome per previously identified genome cluster (i.e. ~species) (Fig. 3). The most recent transitions between the aquatic biomes, if detected, should be represented on the phylogenetic tree as pairs of closest related monophyletic and biome-specific groups. We call these clades monobiomic sister groups (MSGs, Fig. 4a). We identified 310 MSG pairs, with brackish and marine (BM; n=136) and freshwater and brackish (FB; n=119) transitions being twice as numerous as freshwater and marine (FM; n=55), despite the larger number of freshwater and marine MAGs in the pruned tree (Fig. 4b). The number of FB and FM transitions were also significantly lower than would be expected if phylogeny would not correlate with biome annotation (Fig. 4c). This corroborates observations for the previously described genome clustering.

**Fig. 3.**
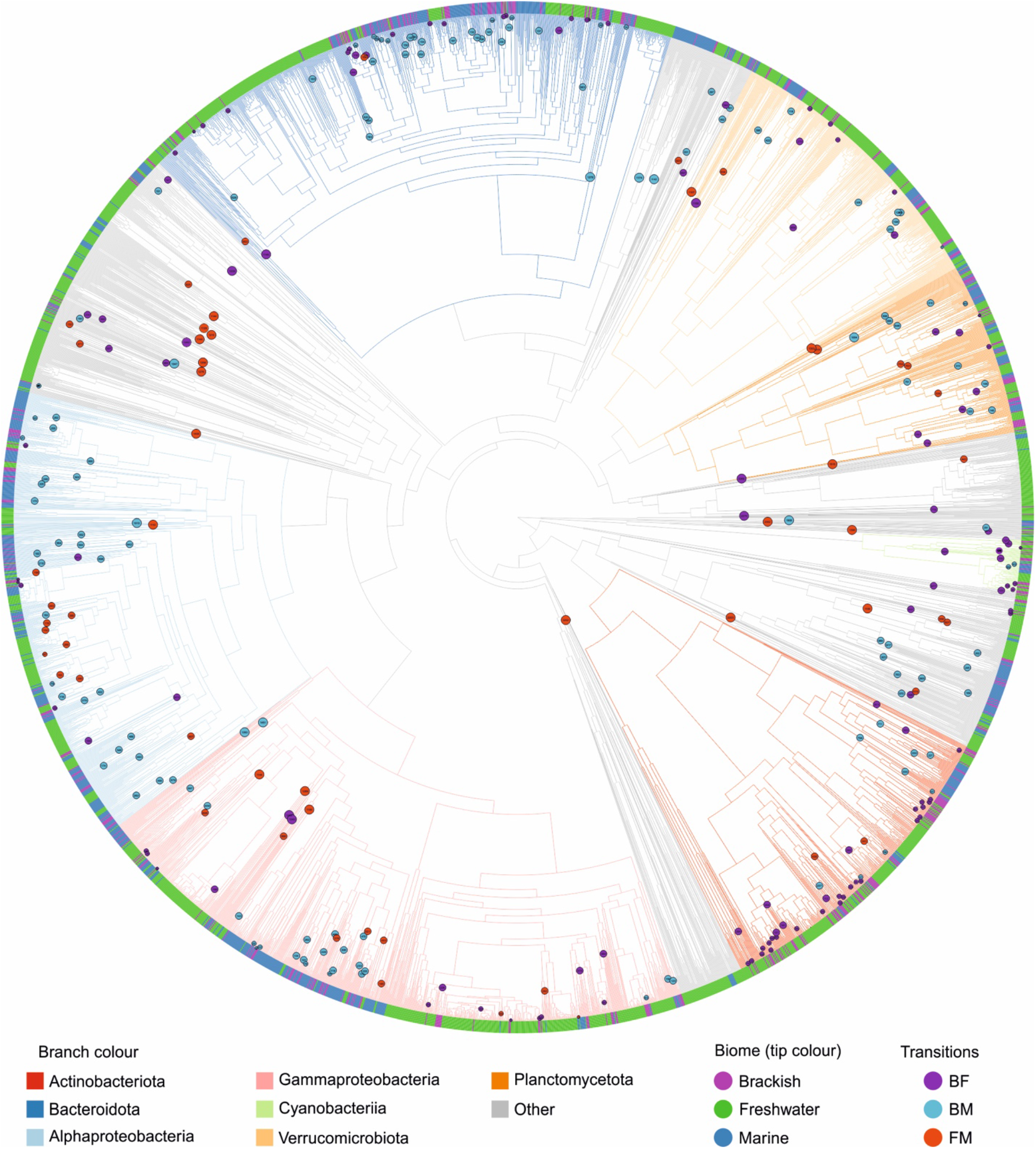
Reconstructed phylogeny of MAGs of the three aquatic biomes. Tree with branch lengths corresponding to estimated minimal times since divergence. Only tips present in the pruned tree are visualized. Identified transitions, i.e. nodes corresponding to most recent common ancestors of MSG pairs, are color-labeled by type.

**Fig. 4.**
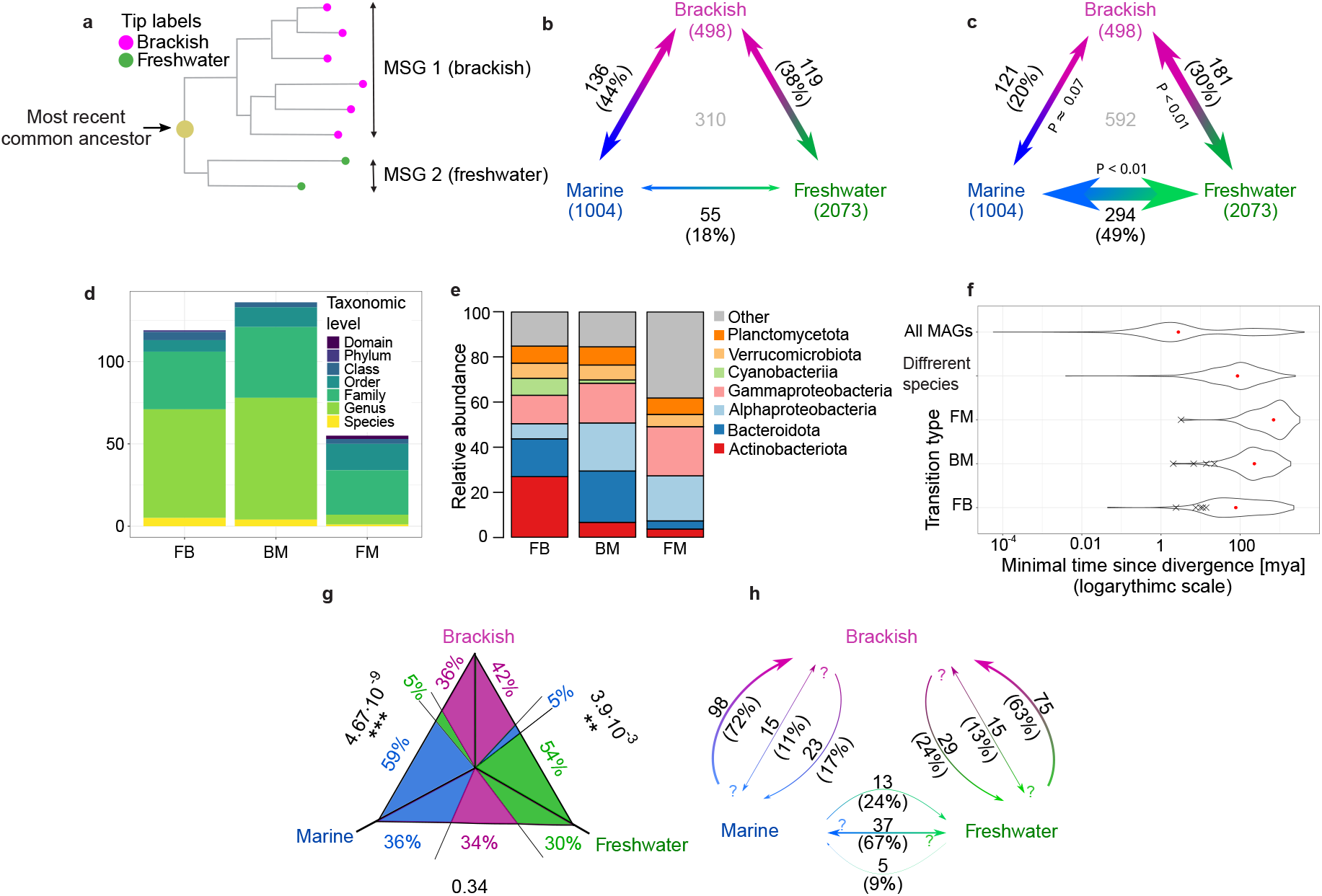
The most recent transitions between biomes. a. Illustration of an MSG pair corresponding to an FB transition **b**. Observed numbers of transitions between the biomes. Numbers of MAGs included in the pruned tree are mentioned in brackets under the biome name. The total number of all identified cross-biome transitions is shown in gray in the middle of the plot. **c.** Expected numbers of transitions between the biomes as predicted by random permutations of biome annotations. On the internal sides of the arrows are given P-values for the estimated numbers being smaller (for FM and FB) or bigger (for BM) than the observed numbers of transitions. **d.** The number of transitions distributed across the lowest shared taxonomic level to which both MSGs could be classified. If no annotation was available at a taxonomic level, a higher one was considered. **e**. Proportion of pairs of MSGs for broader taxonomic groups. **f.** The distributions of estimated minimal times since divergence for all nodes on the unpruned tree (“All MAGs”), nodes corresponding to most recent divergence events (pairs of bacterial species sharing an exclusive MRCA on the pruned tree, i.e. “Different species”), and nodes corresponding to transitions grouped by type. The timing of transitions within >95% ANI clusters is marked as “X”. **g.** Mean likelihoods of ancestral biome-states for transition between the biomes. Next to the sides of the triangles are given p-values for difference in the likelihood for ancestral states of the 2 biomes in which the MSGs for the transition type were identified (for ex. between brackish and freshwater for FB). **h.** Numbers (and proportion) of transitions of each type for which the direction of the transition indicated by the arrows was more probable than the two other ancestral biome-states taken together. Both-sided arrows indicated transitions for which no dominant transition direction was identified.

Most of the identified FB and BM MSG pairs belonged to the same genus (around 55% in both cases), while for FM transitions the lowest-level shared taxonomic annotation was most often (49%) at the family level (Fig. 4d, Supplementary Table S1). Only small proportions (1.8 – 4.2%) of the MSG pairs belonged to the same species (defined either by GTDB annotation or as a >95% ANI genome cluster). The taxonomic distribution of MSG pairs followed the overall distribution in the corresponding biomes for BM and FM, but not for FB, probably due to the overrepresentation of Actinobacteriota and Cyanobacteriia in this transition type (Fig. 4e, Supplementary Fig. S1).

We estimated the minimal time since divergence of the MSG pairs based on phylogenetic distances, using a method which allows for variable evolutionary rates^28^ (Fig. 3, Fig.4f, Supplementary Fig. S2). As time constraints, we used minimal times since divergence of genomes of endosymbiotic bacteria known from fossil records of their hosts^29−37^ (Supplementary Table S2). As obligate endosymbionts have high evolutionary rates that exceed those of free-living bacteria^37^, this model gives conservative estimates of time since the transitions started. The timing differed depending on the transition type (Fig. 4f; *P* < 10^−6^ for all comparisons), FB being the most recent and FM the oldest. The difference in median times since FB and BM transitions could not be attributed to the uneven representation of biomes in the dataset (Supplementary Fig. S2b).

The most recent transition was estimated to have happened at least 44.8 kya, and all other transitions millions of years ago (Supplementary Table S3). Therefore, all the detected transitions started long before the current brackish conditions were established in the Baltic Sea (8 kya^38^) and only the six most recent ones to the period since the Caspian has continuously been brackish (2-3 mya^39^). These results imply that the evolutionary history of the global brackish microbiome^12^ by far predates the formation of these two brackish basins.

To infer the directions of the biome transitions, we conducted ancestral (biome) state reconstructions of the most recent common ancestors (MRCAs) of the MSG pairs, correcting for the uneven representation of biomes in the pruned tree (see Methods). FB and BM transitions were generally more likely to happen into than out of the brackish biome, while for FM transitions no significant direction preference was observed (Fig. 4g-h, Supplementary Fig. S3 and 4).

### Large-scale changes in proteome properties follow cross-biome transitions

We next assessed if changes in proteome properties accompany the biome transitions. The average distributions of isoelectric points (pIs) of the proteins encoded in the genomes were compared between MSGs in a pairwise manner. Significant differences in the proportion of neutral (pI ∈ [5.5, 8.5)) proteins were observed for all the transition types (Fig. 5a-b, Supplementary Data S3), with a higher proportion in the biomes of lower salinity. The frequencies of acidic proteins (pI ∈ [3.0, 5.5)) showed the opposite pattern and were significantly different for the comparisons that included freshwater MSGs, but not for the BM transition type. The results suggest that the higher representation of neutral pIs observed in freshwater bacteria^14^ gradually diminishes with increasing salinity while low proportion of acidic proteins is a distinctive feature of freshwater bacteria. However, the extent of transition-related changes in pI distribution highly depended on taxonomy, with particularly small changes for Actinobacteria (Supplementary Fig. S5).

**Fig. 5.**
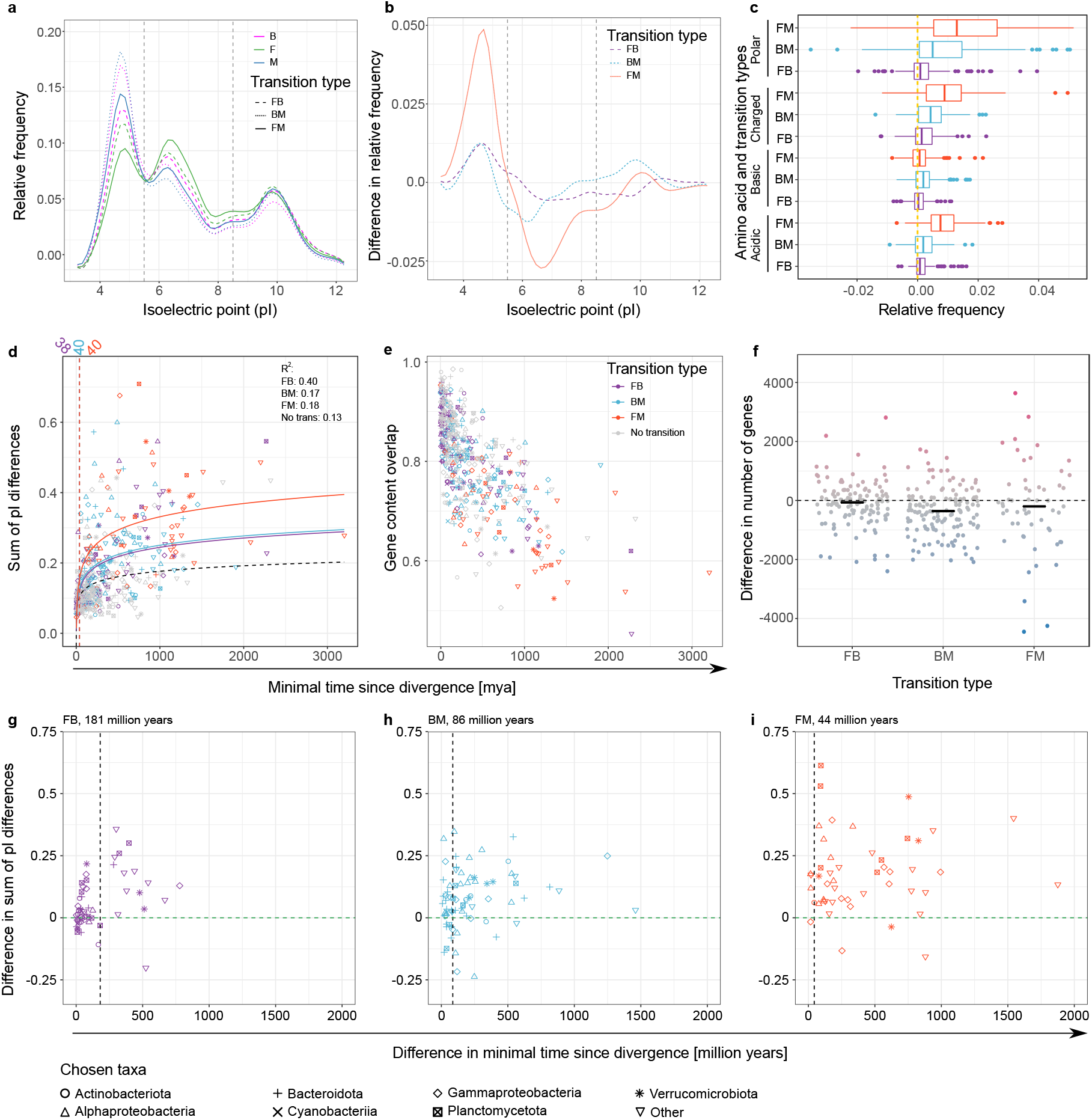
Changes in proteome properties and gene content with cross-biome transitions. a. Averaged distribution of pI values in predicted proteomes of MAGs. The figure is based on the mean frequencies of proteins within 0.5 pH wide bins (from pI ∈ [3.0, 3.5) to pI ∈ [12.0, 12.5)). Vertical lines mark pI values set to distinguish between acidic, neutral, and basic proteins. **b.** Differences in pI frequencies (as in **a.**) across transitions. **c.** Differences in fractions of the proteomes which the chosen categories of amino acids make up. The lower categories are subsets of the higher ones: polar ⊃ (charged = acidic + basic). **d.** Sum of all differences in pI frequency vs. minimal estimated divergence times, for different transition types and for no transition pairs. For no transition (same biome), the most distantly related pairs of monophyletic groups within MSGs were used. A logarithmic relationship was fitted for each transition type and no transition events. The vertical lines and values above correspond to the timepoint when the fit for a transition and no transition events diverge significantly (no overlap in +/− standard error ranges thereafter). **e.** Overlap in gene content vs minimal time since divergence (grouped as in **d**). **f.** Differences in numbers of genes between the pairs of MSGs. The horizontal bars represent median values. **g-i.** Differences in the sum of pI differences between MSG pairs and between the biggest monophyletic groups within the MSGs formed after the transitions vs. differences in minimal times since transition and subsequent within-MSG divergence events for FB (**g**), BM (**h**), and FM (**i**). The vertical lines correspond to the highest values in mya for which the differences of sums of pI differences are not significantly higher than zero. In **b**, **c** and **f**, the value differences are presented as value in the more saline - value in the less saline biome.

Polar, charged, and acidic amino acid frequencies broadly increased with salinity across the MSG pairs (Fig 5c, Supplementary Data S3). Changes were most significant for acidic amino acids, which is a subcategory of the other two categories and may thus also explain the changes observed for those. In contrast, the frequency of basic amino acids differed significantly only for BM transitions.

Among the acidic amino acids, the difference for glutamate was larger and more significant than for aspartate across all the transition types (Supplementary Fig. S6, Supplementary Data S3). For BM transitions, the higher proportion of basic residues in marine proteomes could be attributed to lysine. Big and significant differences in the frequency of alanine across BM and FM transitions suggest that this non-polar amino acid with a relatively “neutral” effect on protein structure^40^ is often substituted by charged residues in transitions to higher salinity. For all transition types, there was significantly less proline in the predicted proteomes at higher salinity.

To investigate the dynamics in how the proteome is reshaped after biome transitions, we compared the differences in pI distributions to inferred times since divergence. The overall change in pI distributions followed different logarithmic relationships for within-MSG (no transition) divergence times as compared to any of the transition types (Fig. 5d). Moreover, the differences in pI distribution across transitions were consistently larger than changes resulting from within-MSG (no transition) divergence events when the latter were at least 44 to 181 million years more recent (Fig. 5g-i). This suggests that proteomes are under strong selection and continue to evolve to match the new salinity conditions long after the switch to a new biome (Supplementary Fig. S7).

### Specific changes in gene content accompany cross-biome transitions

For each transition type, we were able to identify specific functional genes (KEGG orthology groups [KOs]), significantly over-represented in the MSGs of one of the compared biomes after correcting for differences in estimated completeness of the MAGs (Fig. 6, Supplementary Data S4). However, transitions did not change the gain and loss rates (Fig 5e), even though gene numbers significantly differed for BM transitions (Fig. 5f, Supplementary Data S3), with brackish MAGs having a median of 386 more genes. Therefore, the gains or losses of the same gene functions (KOs) we observed in multiple independent transition events most likely result from analogous changes in selective pressures caused by the biome transitions.

**Fig. 6.**
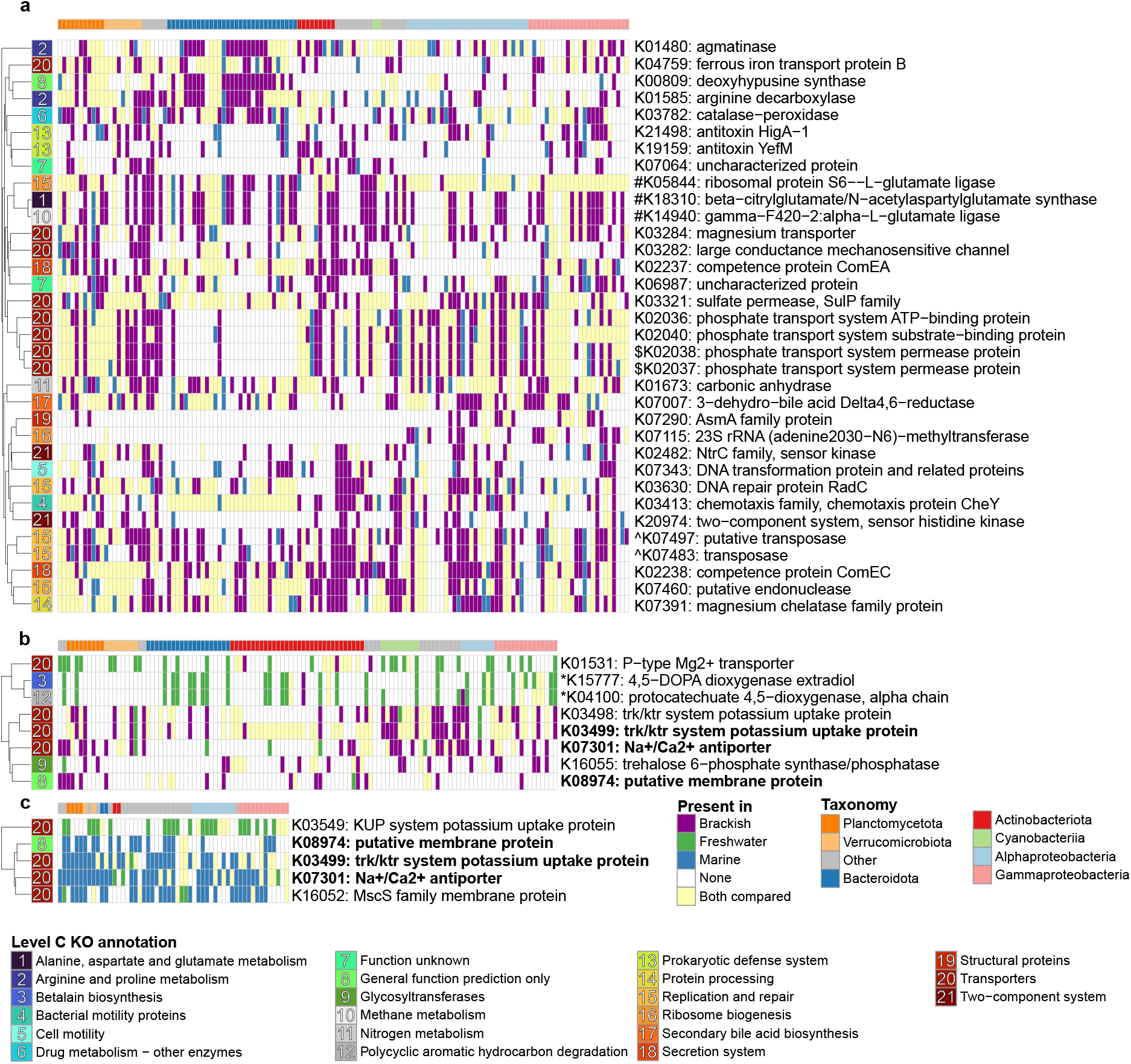
Functional genes differentially present across transitions (“gain-loss maps”) Genes present in only one MSG from BM (**a**), FB (**b**), and FM (**c**) pairs of MSGs. Each column is a pair of MSGs, with taxonomic annotation marked above, and each row is an orthologous group, with its KO number and a descriptive name given on the right and a broader (KO level C) functional annotation marked to the left. To correct for labels inconsistent with the literature reference, the annotations of K01531^45^ and K07301^96^ were manually curated. The category “Enzymes with EC numbers” (represented by K00809) has been merged with “General function prediction only”. The genes are marked as present in an MSG if they were found in >0.5 of the MAGs of the MSG. Only KOs found significantly (FDR adjusted *P* < 0.1) differentially present across transitions in a pairwise analysis are displayed. Descriptions of genes identified in more than one transition type were marked in bold. Groups of genes with at least one gene being in >50% of the cases being annotated also as the other(s) (Supplementary Data S4) were marked by adding #, $, ^ or * before the KO number.

Transporters were the only functional category (KO level C) over-represented in all transition types (Fig. 6, Supplementary Figure S8), some with known roles in adaptation to altered salinity. For FB transitions, brackish MAGs more frequently contained components of two (or of the same^41^) potassium uptake systems (K03498 and K03499) co-acting to allow growth in higher osmotic pressure^42^. A component of the uptake system with higher affinity, K03499^42^, was also more prevalent in marine genomes across FM transitions. Additionally, for both FB and FM MSG pairs, a Na^+^/Ca^2+^ antiporter (K07301) was more often found in the more saline biomes. Analogous changes in presence of the same K^+^ uptake systems and of Na^+^/Ca^2+^ antiporters were reported in a comparison of terrestrial and marine *Flavobacteriaceae^43^*. Finally, K03282, more frequently present in brackish across BM transitions, is a mechanoreceptor involved in managing hypoosmotic stress^44^.

Two distinct magnesium uptake systems^45,46^ were more frequently present in lower-salinity biomes for FB and BM MSG pairs (K01531 and K03284, respectively). Moreover, for BM transitions, brackish MAGs displayed a higher presence of chemotaxis (K03413) and two-component system sensors (K02482 and K20974) involved in regulation of motility^47,48^. While Mg^2+^ is a major component of sea salt^11^ and salinity can alter chemotactic responses and motility^49–51^, to the best of our knowledge, there are no previous reports on the eco-evolutionary relevance of these genes. For FB, a gene connected to either pigment synthesis (K15777) or aromatic hydrocarbon degradation (K04100) was more often found in freshwater MAGs. Both of these functions may be connected to salinity-related changes in dissolved organic carbon concentration (see Supplementary Discussion for more details).

Other differentially present genes were less directly connected to physicochemical differences between the biomes, suggesting more complex and/or cryptic biological mechanisms associated with transitions. For BM transitions, genes connected to various metabolic pathways (including modifications of charged amino acids), as well as transposases, antitoxins, and competence proteins, were more often present in brackish MAGs than their marine counterparts. Typical housekeeping genes were absent among the differentially present genes, which verifies the specificity of the method.

## Discussion

In this study, we combined several large-scale datasets, which allowed us to comparatively analyze bacterial genomes from freshwater, brackish and marine biomes at an unprecedented scale. Including brackish genomes allowed for a more detailed analysis of the evolutionary history of aquatic bacteria than in previous studies focusing on the two biomes at the extremes of the spectrum^9–11,16,43, 52, 53^. The brackish biome is a significant part of our global waters and combined, the two major brackish systems analyzed here have a similar volume (~99,000 km^3 54^) as all freshwater lakes on earth (≤102,000 km^3 55^), albeit only a fraction of the volume of the global ocean (~1.35 billion km^3 56^).

We observed bacterial communities from the different biomes to be separated on at least the species level - nearly all >95% ANI genome clusters were restricted to a single biome. Brackish genome clusters often had representatives from both the Baltic and Caspian Sea. These results provide further support that the brackish biome hosts a distinct, globally distributed set of bacterial species^12^. However, the fact that populations of individual species were more genetically differentiated between the two brackish seas than across the strong salinity gradient within the Baltic Sea indicates that geographical barriers still have a structuring effect on the distribution of aquatic bacteria. This implies a limitation in microbial dispersal. While it might be of limited relevance on a large evolutionary timescale, it is likely to be of crucial importance when an ecosystem is disturbed, as it may take substantial amounts of time for migrating strains to fill the niches which open up as a consequence of the disturbance.

Alternatively, local bacteria could adapt to environmental changes, maintaining a diverse and functioning ecosystem throughout the process. This is unlikely to be the case when it comes to changes in salinity spanning across biome barriers however, since neither the formation of the Baltic nor the Caspian Sea caused an observable increase in transition rates, which is consistent with previous notions of major geological events having little impact on bacterial speciation rates^57^. Therefore, most of the local microdiversity which we uncovered through population genomics is likely to be lost as a consequence of such environmental changes. Ultimately, the destruction and disturbance of environments, enhanced in the times of global change, not only threaten local microbial diversity in the short run but also diminish genetic diversity even within global species.

Our analyses support previous concepts of freshwater-marine transitions being rare^9–11^ and show more common but still rare freshwater-brackish and brackish-marine transitions, which most often are directed towards the brackish biome. However, as the three biomes have continuously existed throughout geological history and bacterial species appear to be globally distributed within each of them^2^, one might ask why any recent transitions have even happened. A plausible scenario may be that genetic innovation (gene gain/loss or mutation) can make a lineage competitive in a different biome, either by opening up a new niche or out-competing others in an existing one, in accordance with the stable ecotype model^58^. However, the benefits of the innovation need to outweigh the decrease in fitness due to switching to a new salinity regime. Thus, the smaller salinity difference between the brackish and the other biomes would explain why it is associated with more transitions.

The more frequent transitions into than out of the brackish environment are harder to explain. For FB, the reason could simply be hydrological, i.e. that freshwater usually flows into brackish water rather than the other way around. For BM, differences in population sizes may instead be the explanation, as both the volume and surface of the ocean are much bigger than of all brackish waters. Beneficial mutations are thus more likely to appear among the more numerous marine bacteria. We also observed that there was a small fraction of FB and BM transitions with unusually high third-biome ancestral states (Supplementary Fig. S4). These events might correspond to FM transitions through brackish waters, where an intermediate period in brackish water would allow for a more gradual adaptation than between FM directly.

Our results corroborate previous reports on differences in predicted proteomes at different salinities^14^, and support them with a systematic analysis across the bacterial tree of life. These changes likely occur over long evolutionary timescales after the actual transition, meaning that in the period after the transitions, bacteria may have a suboptimal protein repertoire in their new environment. This environmental mismatch would potentially lower the fitness of newly transitioned lineages in addition to the bottleneck effects, traces of which are present in LD12 clade genomes^59^, and which can lead to fixation of slightly deleterious mutations^60^.

We observed a shift towards more acidic protein pIs at higher salinity, also reported before^14,61^, and showed the underlying proteome-scale increase in proportion of acidic amino acids, most notably glutamate. This likely reflects that charged amino acids increase protein solubility at higher ionic strength^62,63^. However, an even stronger pattern observed was the proportion of neutral proteins gradually diminishing with increasing salinity, which is also known from previous studies but lacks explanation^14^. We hypothesize that this effect might be due to diminishing extracellular pH variation as the concentration of buffering salts increases^11,64^. Moreover, Na^+^-coupled transport is the key mechanism of pH homeostasis under alkaline conditions^65^, to which both freshwater^11^ and brackish environments^64,66^ transcend. Consistently, lower NaCl concentrations have been connected with higher intracellular pH variation in alkaline conditions^65^. Thus, neutral proteins could be soluble and active outside as well as inside the cell under pH variations in less saline environments. Unlike freshwater, brackish pH rarely changes to an acidic state^64,66^, which may explain why the difference in frequency of acidic proteins was less significant for BM than for other transition types, accompanied by a change in the frequency of not only acidic but also basic amino acids.

For BM transitions, we identified genes involved in glutamate post-translational modifications (K03412, K05844, K14940, K18310) to be more frequent in brackish genomes. All these modifications increase the acidity of residues^67–70^. Thus, depending on the specificity of the enzymes, brackish bacteria may change the charges (and pIs) of their proteins to adjust to different salinities (and varying pH). We also observed that brackish bacteria were enriched in gene functions for other physiological responses to environmental cues, including chemotaxis, transcriptional regulation, as well as synthesis of trehalose and polyamines (both osmolytes) (see Supplementary Discussion). Thus, phenotypic plasticity might be an important hallmark of brackish bacteria, sustaining the populations across the physicochemical gradients typical for many brackish environments.

BM transitions seem to expose bacteria to a different set of mobile genetic elements (MGEs). This is suggested by a set of transposases (previously reported to be unusually abundant in the Baltic Sea^71^), competence proteins, and, indirectly, antitoxins^72^ being more often present in the brackish MAGs (Fig. 6a). Consequently, more MGEs in the brackish MAGs could explain the higher numbers of genes as compared to the marine relatives (Fig. 5f). MGEs tend to increase gene gain and loss processes^73,74^, and may thus facilitate fast adaptation to new physiochemical conditions. It also promotes the establishment of a recombination barrier between biomes^75^. As homologous recombination inhibits divergence^75–79^, these effects would give a further mechanistic explanation for the “species-level” separation.

We used a novel comparative genomics approach to leverage the large phylogenetic scale of the analysis. It allows systematic comparison of close relatives differing in categorical states and evolutionary dynamics of transitions between these states. Defining the transitions as pairs of MSGs, we could meaningfully classify them without relying on often ambiguous ancestral state inference (which we still perform, but only for inferring directions of transitions and in the process correcting for imbalances in the dataset in an evolutionary history-informed manner). Moreover, comparative genomics and evolutionary dynamics can be linked to each other with this approach. That can be instrumental to move beyond mere correlations of ecological and genomic changes toward their causal relationship, as we showed by unraveling the prolonged nature of post-transition proteome reorganization. While correlation-based methods can also be adjusted for a taxonomically balanced comparative genomics analysis ^80^, it is much harder to connect these results to specific evolutionary events. Furthermore, with the new framework, we could correct for varying levels of genome completeness across MSG pairs, and not downsampling all the MAGs to the same, lowest, level, therefore minimizing the information loss. Consequently, we detected gains or losses of genes obviously related to adaptation to salinity but not any housekeeping genes. We were thus able to distill the traces of repeated gene gains and losses due to habitat change from the taxonomic effects which govern the differences in gene content of whole metagenomes ^16^. At the same time, we identified convergent changes connected with cross-biome transitions in diverse bacterial lineages. These results, and the approach, can serve as a basis to disentangle the effects of transitions from consequences of parallel events and/or random changes when studying evolutionary histories of specific taxa.

Ultimately, we extend the view of transitions between aquatic biomes and highlight the brackish environment, showing trends specific for certain transition types and common across all of them. We report patterns prevalent across the bacterial tree of life, providing supplementary materials that allow investigation of the changes for chosen taxonomic groups. We observed many changes related to previously undescribed adaptation mechanisms and/or poorly characterized genes, both of which call for further exploration. In the future, phylogenomic analyses should be supplemented with experimental and ecological approaches. This would allow examining the relevance of the unraveled adaptation mechanisms in the context of ecosystem functioning and physiology.

## Methods

### MAG collection and sources

Representative metagenome-assembled genomes (MAGs) from freshwater, brackish, and marine biomes were collected from seven previous studies as specified in Table 1. Freshwater MAGs originate from two North American lakes (Lake Mendota and Trout Bog), 44 lakes of the StratfreshDB (mainly in Sweden, Finland and Canada), and Lake Baikal (Russia)^18–20^. Brackish MAGs originate from the Baltic Sea^17^ and the Caspian Sea^21^. Finally, marine MAGs that were published by Tully et al.^22^ and Delmont et al.^23^ originate from the Tara Oceans Survey^81,82^.

### MAG statistics, ANI calculations and classification

Completeness and contamination of the collected MAGs were assessed using CheckM^83^. MAGs with ≥75% completeness and ≤5% contamination were considered for further analyses. Pairwise Average Nucleotide Identity (ANI) between all MAGs was calculated using FastANI (V1.2) and MAGs were clustered at the threshold of 95% ^84^. Clustering of MAGs was done using hclust and cuttree functions in Rstudio. To remove redundant information and reduce the complexity of the phylogenetic tree we kept one random MAG per cluster per biome to generate a pruned tree. Taxonomic affiliation of MAGs was assigned using the “classify_wf” workflow of the Genome Taxonomy Database Toolkit (GTDB-tk) (v 0.3.2)^25,27^. To compare proportion of inter-biome and brackish inter-basin clusters, we used Fisher’s exact test conditioned on the lower number of clusters (2 categories: the basin/biome with a lower number of clusters, and the number of shared clusters).

### Population genomics

Population genomics was conducted with the POGENOM software as described in Sjöqvist et al.^26^. The same set of 22 MAGs (from^17^) were used as references for mapping of metagenomic data from the Baltic^17^ and Caspian^21^ seas. Only five ese obtained sufficient coverage in the three Caspian Sea samples and were included in the downstream analysis. The five MAGs all represent different (>95% ANI) genome clusters. The Input_POGENOM pipeline was used for the automatic generation of input files for POGENOM., The pairwise *F_ST_* values output from POGENOM were used to conduct Principal Coordinates Analysis (PCoA) using the Vegan package in *R*. Maps were plotted with the rworldmap package. For parameter settings of Input_POGENOM and POGENOM, see Sjöqvist et al.^26^.

### Phylogeny reconstruction

The “de_novo_wf” function of GTDB-tk (v 0.3.2) was used to reconstruct the phylogeny of MAGs^25,27^. The WAG (Whelan and Goldman) amino acid evolution model was selected in the infer step and phylum Patescibacteria was selected as the outgroup^85^. Most downstream phylogenetic analyses were performed in RStudio (V3.6.2) with the R packages ape, phytools, phangorn and ggplot. The multi2di function in R was used to resolve multichotomies (that appear because some MAGs were nearly identical) in the phylogenetic tree. Since this function rendered some branches with zero length, we arbitrarily changed all zero-length branches to 5 – 10^−4^ (which was the smallest length of all other branches) to ensure all branch lengths were > 0.

### Gene prediction and annotation

Prodigal (v2.6.3) was run with default settings to identify protein-coding genes^86^. Gene function annotation was performed with eggNOG-mapper (v2.0.0), based on the eggNOG v5.0 database^87,88^. A custom Python script was used to count the number of occurrences of each Kegg Ortholog (KO) in each annotated MAG.

### Identifying monobiomic sister groups (MSGs) for comparative analyses

Pairs of MSGs (Fig. 4a) were identified based on the pruned tree. The most recent common ancestors (MRCAs) of the monobiomic groups were identified by choosing nodes for which on each of the two downstream branches there were MAGs from a different and only one biome. Transition types were defined by the two biomes from which the MAGs originate and do not take into consideration inferred direction of the transition. The biggest possible pairs of monophyletic groups within MSGs (coming from the same biome; only one pair for each MSG with >1 representative MAG on the pruned tree) were used as “no transition” divergence events.

### Expected number of transitions

Biome annotation of the tips of the pruned tree was randomly permuted 100 times and MSG pairs were re-identified using the same tree structure and the random biome annotations. Rounded averages of transitions obtained in all the iterations were used as the expected numbers. One-sided P-values were calculated as the number of iterations in which a lower/higher number of transitions of a type was observed. The fractions of each transition type among all expected transitions were used as probabilities for *X*-squared test on the observed values.

### Minimal time since divergence

Six divergence-time constraints were set as minimal estimates of time since divergence for pairs of obligate endosymbiont taxa based on fossil record of host species^29–37^ (Supplementary Table S2, Supplementary Data S2). Divergence times were calculated using RelTime^28^ based on the unpruned trees, including GTDB reference sequences. Maximum relative rate ratio was set to 100. To correct for the difference between the placing of the root based on phylogenetic distances only^85^ and the most probable position of the last bacterial common ancestor (LBCA) on the tree^89^, *Fusobacteriota* were chosen as an outgroup. Only the tips present in the pruned tree were chosen for further analyses.

### Transition directions

One MAG from each MSG was randomly chosen. Additional MAGs from outside from the MSGs were randomly chosen to downsample the dataset to the same number of genomes from each biome, equal to the lowest number for any of the biomes (396 for brackish after removing multiple

MAGs within each MSG). “ace” function from the “ape” package in R was used to obtain maximum likelihood ancestral states of the MRCAs of MSG pairs using an all-rates-different model^90^. Pairwise Wilcoxon rank-sum test of likelihoods of the ancestral states for the two biomes between which transition occurred was performed for each transition type. Random selection of MAGs within and outside the MSGs was iterated 100 times and mean likelihoods and p-values were used for further statistical analyses.

### Comparison of isoelectric points and amino-acid frequencies

Isoelectric points (pI) of all proteins within inferred bacterial proteomes were obtained using the pepstats function from the EMBOSS package (v6.6.0)^91^. To get amino acid and amino acid category frequencies the whole proteomes were concatenated into one sequence before running separate pepstats analyses. pI frequencies within bins of pH width of 0.5 spanning the range from 3.0 to 12.5 were used to average, plot and compare the distributions of pIs between MSGs. Average values over MAGs (95% ANI cluster representatives) were used for MSGs consisting of more than 1 MAG.

For analysis of total changes in pI distribution, the absolute values of differences for each 0.5 pH wide bin were summed up. To compare transitions to divergence events within the same biome, within each MSG bigger than 1 representative MAG, a pair of monophyletic groups which diverged the longest time ago were identified. The differences between these “no transtion” (within-MSG) monophyletic groups were compared with differences across the ancestral nodes (MRCAs of MSG pairs) in a pairwise manner.

### Functional gene overlap and gene numbers

Occurrences of KOs were transformed into a binary table of gene (KO) presence/absence. Gene overlap was calculated as: 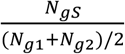, where *N_gs_* is the number of KOs shared between 2 genomes and *N_g1_* and *N_g2_* are the total numbers of KOs present in the respective genomes^75^. Formulas *log(gene overlap) ~ divergence times* and *gene overlap ~ divergence times* were used to fit logarithmic and linear models for each transition type, including no transition (see Comparison of isoelectric points and amino-acid frequencies). The models were compared using a method based on the least-square means statistic^92^. Numbers of distinct polypeptide sequences in the predicted proteomes were used as the gene numbers.

### Gene presence/absence analysis

Occurrences of KOs were transformed into a binary table of gene (KO) presence/absence. Random pairs of MAGs from the MSGs were chosen for comparison for each transition type. To correct for differences in genome completeness, the presence count for the more complete MAG in each pair was randomly downsampled to the level of the less complete MAG. For each transition type, KOs present in at least one of the MAGs within any of the MSGs were analyzed. For each of the KOs, differences in presence between MAGs from each combination of two biomes were analyzed using pairwise Wilcoxon rank-sum test^93^. The procedure was iterated 100 times, starting from the step of randomly choosing pairs of MAGs. The Benjamini-Hochberg method^94^ was used to obtain false-discovery rate (FDR)-adjusted p-values (q-values), and average q-values from the 100 iterations for each KO were used to determine the significance of difference in each KO’s presence/absence. The identified KOs were annotated using KEGG orthology^95^ table collected from the KEGG website on the 24th of November 2021.

### Statistical analyses

If not stated otherwise, pairwise Wilcoxon rank-sum test^93^ was used to compare values across MSG pairs of a certain type. If not stated otherwise, family-wise error rate (FWER) ≤ 0.05 was considered as the threshold of significance. The analyses were performed using R.

## Data availability

All the MAGs used in this study are made publicly available via their original publications as referenced in the text and Table 1. Data files describing intermediate data analysis steps, including the phylogenetic trees, and the code for in-house data transformation procedures, are available at https://doi.org/10.17044/scilifelab.20732170.

## Funding

This study was supported by a grant from the Swedish Research Council VR to A.F.A. (grant no. 2021-05563). Publication funding was provided by KTH Royal Institute of Technology.

## Author contributions

A.F.A designed the study in consultation with all coauthors. K.T.J., M.M, A.F.A, Z.D., F.D. analyzed the data. K.T.J. and M.M. wrote the first draft of the manuscript. All authors contributed to the interpretation of the data, revised the manuscript and approved the final version before submission.

## Acknowledgement

Authors would like to thank Dr. Moritz Buck for providing access to the StratFreshDB MAGs that were included in this study.

## Conflict of interest

Authors declare no conflict of interest.

## Notes

### Competing Interest Statement

The authors have declared no competing interest.

### Summary of Updates

Updated discussion

