## Supplementary Information for "Large-scale phylogenomics of aquatic bacteria reveal molecular mechanisms for adaptation to salinity"

Supplementary Data Description…………………………………………………………. 2
Supplementary Tables …..,…………………………………………………………….  3
Supplementary Figures …….…………………………………………………………..  6
Supplementary Discussion……………………………………………………………. 14

### Supplementary Data Description

#### Supplementary Data 1

A spreadsheet with detailed results of clustering and MSG identification. Contains a table annotating MAGs to >95% ANI clusters and the representatives chosen for further analysis marked as well as sheets with just the representatives, clusters common between the brackish basins and between the biomes. The 2$^{nd}$ sheet (MSG_table) is a table with all the MAGs within identified monobiomic sister groups (MSGs), annotated to appropriate transition_ID, biome and transition type. Taxonomic classification and transition times and directions are also included in this table.

#### Supplementary Data 2

Constraint file used for estimating time since divergence, input for RelTime (MEGA11). Minimal estimates of time since host species diverged [mya], based on the fossil record, were used to set the constraints (see Supplementary Table S1).

#### Supplementary Data 3

A spreadsheet with detailed results of comparison of isoelectric point (pI) distributions and amino acid compositions of proteomes across pairs of MSGs (transitions). Statistics (p-values and differences sizes) for pairwise comparisons of inferred proteome properties and composition, i.e. i) relative frequencies of acidic, neutral and basic (pI categories) proteins; ii) genome sizes as defined by number of inferred protein-coding genes; iii) amino acid relative frequencies; iv) relative frequencies of amino acids categories. Each set of statistics is followed by a table with changes across each of the identified transitions (MSG pairs) given separately, connected to transition ID, taxonomy, and transition type ("tr_diffs" in name of the sheet; for pIs, changes for the 3 protein categories as well as 0.5 pH are given, the latter named by the value in the middle of the range).

#### Supplementary Data 4

A spreadsheet with detailed results on identified significantly differentially present (gained/lost) genes across pairs of MSGs (transitions). Sheets 1-3: tables with all the significant (FDR < 0.1, shaded in orange) differentially present genes. For FB and FM type transitions additional genes were added to the table to show at least the top 25 most significant genes regardless of the FDR values. Sheets 4-6: Biome(s) in which the differentially present KOs were found across the identified transitions (MSG pairs), i.e. the data presented in Fig. 6 in text form and annotated to more specific taxa and single transition events. Sheets 7-9: Fraction of cases in which gene A (row) was also annotated as gene B (column), based on {transition type}.annotation.gz files.

### Supplementary Tables

**Supplementary Table S1 |** The number (proportion) of transitions of each kind for which a given taxonomic level is the lowest to which both MSGs belong. If no annotation was available at a taxonomic level, a higher one was considered. Pairs of MSGs belonged to the same >95% ANI cluster, but were not annotated to the same species by GTDB (3 FB cases), the lowest taxonomic level was considered to be species.

| | **FB** | **BM** | **FM** |
|---|---|---|---|
| **Domain** | 0 (0%) | 0 (0%) | 2 (3.6%) |
| **Phylum** | 1 (0.84%) | 0 (0%) | 0 (0%) |
| **Class** | 5 (4.2%) | 3 (2.2%) | 3 (5.5%) |
| **Order** | 7 (5.9%) | 12 (8.8%) | 16 (29%) |
| **Family** | 35 (29%) | 43 (32%) | 27 (49%) |
| **Genus** | 66 (55%) | 74 (54%) | 6 (11%) |
| **Species** | 5 (4.2%) | 4 (2.9%) | 1 (1.8%) |

**Supplementary Table S2 |** Constraints used to estimate minimal time since divergence (as in input constraint file for RelTime Supplementary Data S2, used by the tool to find MRCAs of given bacterial species and set provided divergence times on these nodes). The choice of species pairs to include was based on Kuo and Ochman 2009. Minimal estimates of time since host species diverged, based on the fossil record, were used to set the constraints. mya - million years ago.

| **Endosymbiont** | **Hosts** | **Time since divergence [mya]** | **Sources** |
|---|---|---|---|
| *Buchnera aphidicola* str. Sg and str. Ua | *Schizaphis graminum* and *Uroleucon ambrosiae* | 50 | Moran et al. 1993; Kim, Lee, and Jang 2011 |
| *Buchnera aphidicola* str. Sg and str. Sc | *Schizaphis graminum* and *Schlechtendalia chinensis* | 80 | |
| *Buchnera aphidicola* and *Wigglesworthia glossinidia* | *Schizaphis graminum* and *Glossina morsitans morsitans* | 120 | |
| *Wigglesworthia glossinidia* | *Glossina brevipalpis* and *Glossina morsitans morsitans* | 23 | Cockerell 1907; Thao et al. 2000 |
| *Blattabacterium* sp. str. Cpu and str. MADAR | *Cryptocercus punctulatus* and *Mastotermes darwiniensis* | 130 | Lo 2003 |
| *Blattabacterium* sp. str. Cpu and *Candidatus Sulcia muelleri* | *Cryptocercus punctulatus* and various sap-feeding insects | 260 | Shcherbakov 2002; Moran, Tran, and Gerardo 2005 |

**Supplementary Table S3 |** Top 10 most recent estimated minimal times since (the start of) transition. Transition IDs as in Supplementary Data S1

| **Transition ID** | **Minimal time since the start of transition [mya]** | **Transition type** | **Taxon** | **Basin of origin of the brackish MSG** | **Is the transition within a >95% ANI cluster?** |
|---|---|---|---|---|---|
| T_20 | 0.0448 | FB | Cyanobacteriia | Baltic | No |
| T_181 | 1.82 | FB | Planctomycetota | Caspian | No |
| T_277 | 2.04 | BM | Gammaproteobacteria | Caspian | Yes |
| T_234 | 2.40 | FB | Alphaproteobacteria | Baltic | Yes |
| T_106 | 2.46 | FB | Bacteroidota | Baltic | No |
| T_304 | 3.22 | FM | Gammaproteobacteria | NA | Yes |
| T_267 | 5.41 | FB | Gammaproteobacteria | Baltic | No |
| T_233 | 5.81 | FB | Alphaproteobacteria | Baltic | No |
| T_30 | 6.70 | BM | Verrucomicrobiota | Baltic | Yes |
| T_300 | 7.08 | FB | Gammaproteobacteria | Baltic | No |

### Supplementary Figures

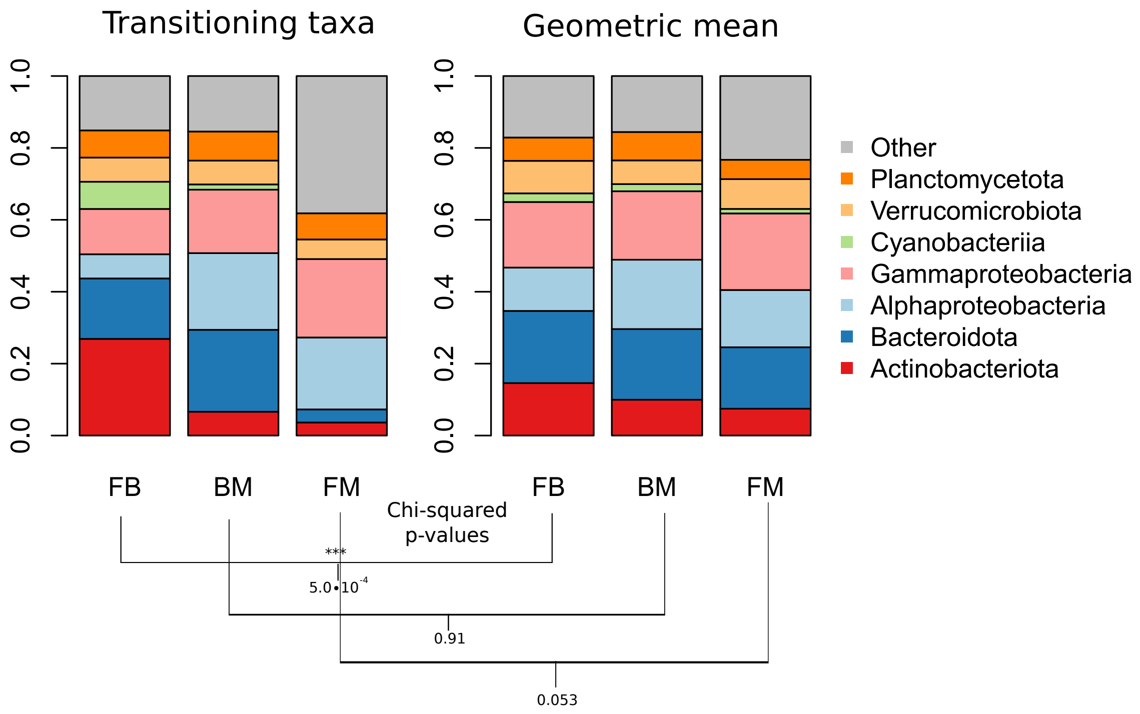


**Supplementary Fig. S1 | Observed proportion of MSG pairs annotated to each of the chosen taxonomic groups vs expected numbers** (geometric means of the proportion of lineages from each of taxonomic groups in respective biomes between which inferred transitions happened). *P*-values obtained from *X*-squared are given for each transition type below the bar charts.

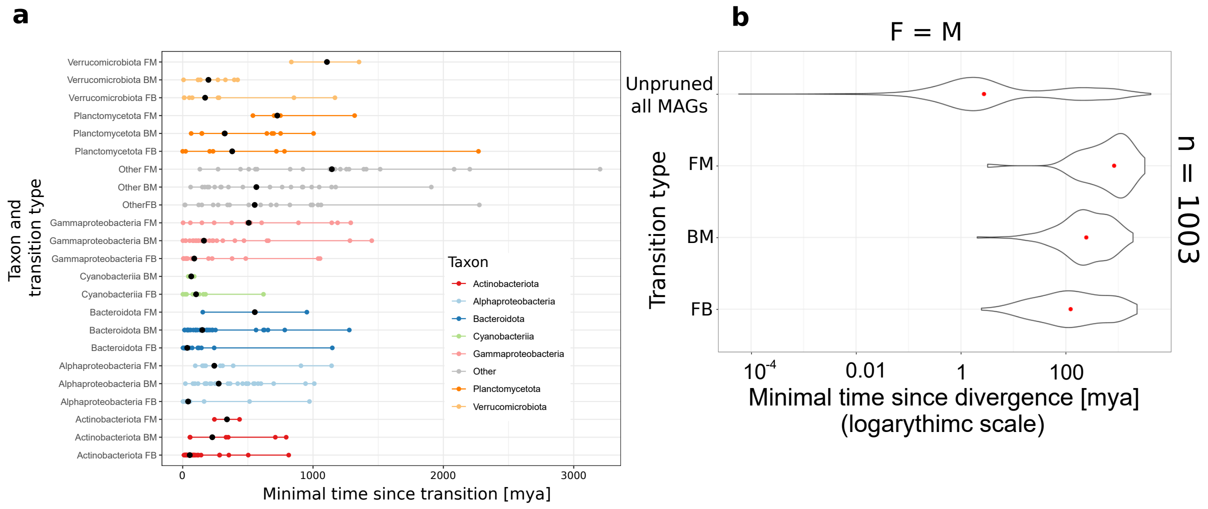


**Supplementary Fig. S2 | Estimated times since the start of transitions a.** grouped by taxa and **b.** based on the pruned tree further subsampled to the same number of freshwater and marine species. **a.** Dots correspond to estimated times since the start of transitions, and horizontal lines connect the most recent and the most ancient ones for each taxonomic group & transition type combination present in the data. **b.** The distributions of estimated minimal times since divergence for all nodes on the unpruned tree ("All MAGs") and for transitions of each type based on the subsampled tree.

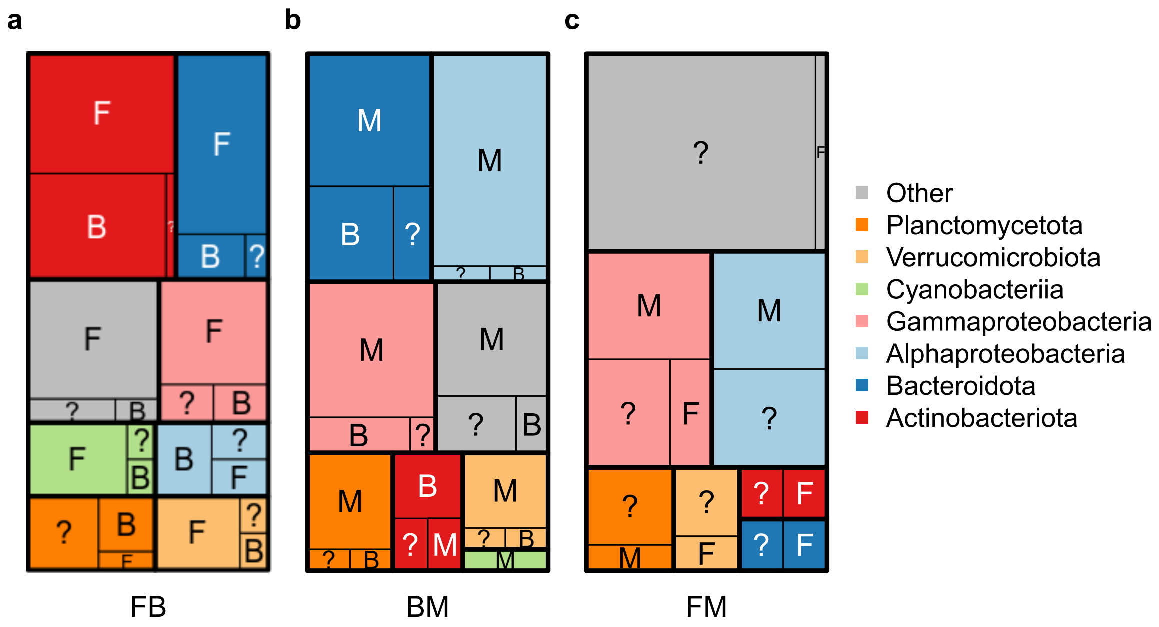


**Supplementary Fig. S3 | Treemaps of dominant transition directions within taxa** for **a.** FB, **b.** BM and **c.** FM transitions. The sizes of the boxes correspond to numbers of transitions for which the biome-ancestral state indicated by the letter (F - freshwater, B - brackish, M - marine) had higher likelihood than the two other ancestral biome-states taken together (*i.e.* likelihood >0.5). "?" indicates that the likelihood of all the ancestral states was <0.5.

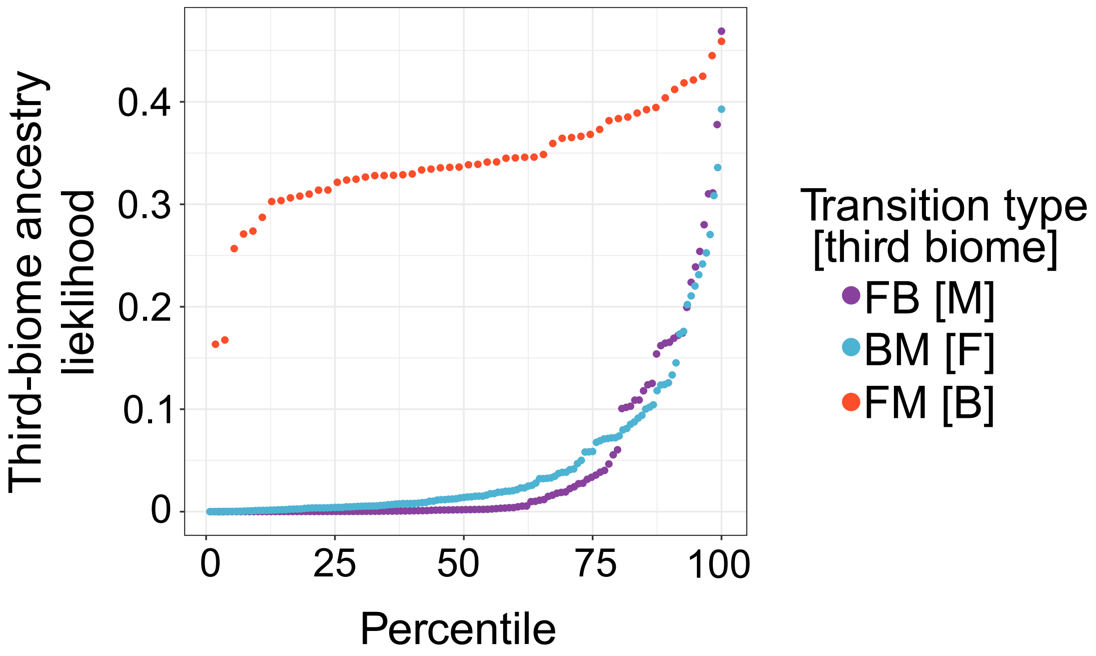


**Supplementary Fig. S4 | The likelihood of MRCAs of MSG pairs originating from a different biome than any of its descendant MAGs (i.e. having a third-biome ancestral state).** The transitions are ordered by increasing likelihood of the third-biome ancestral state.

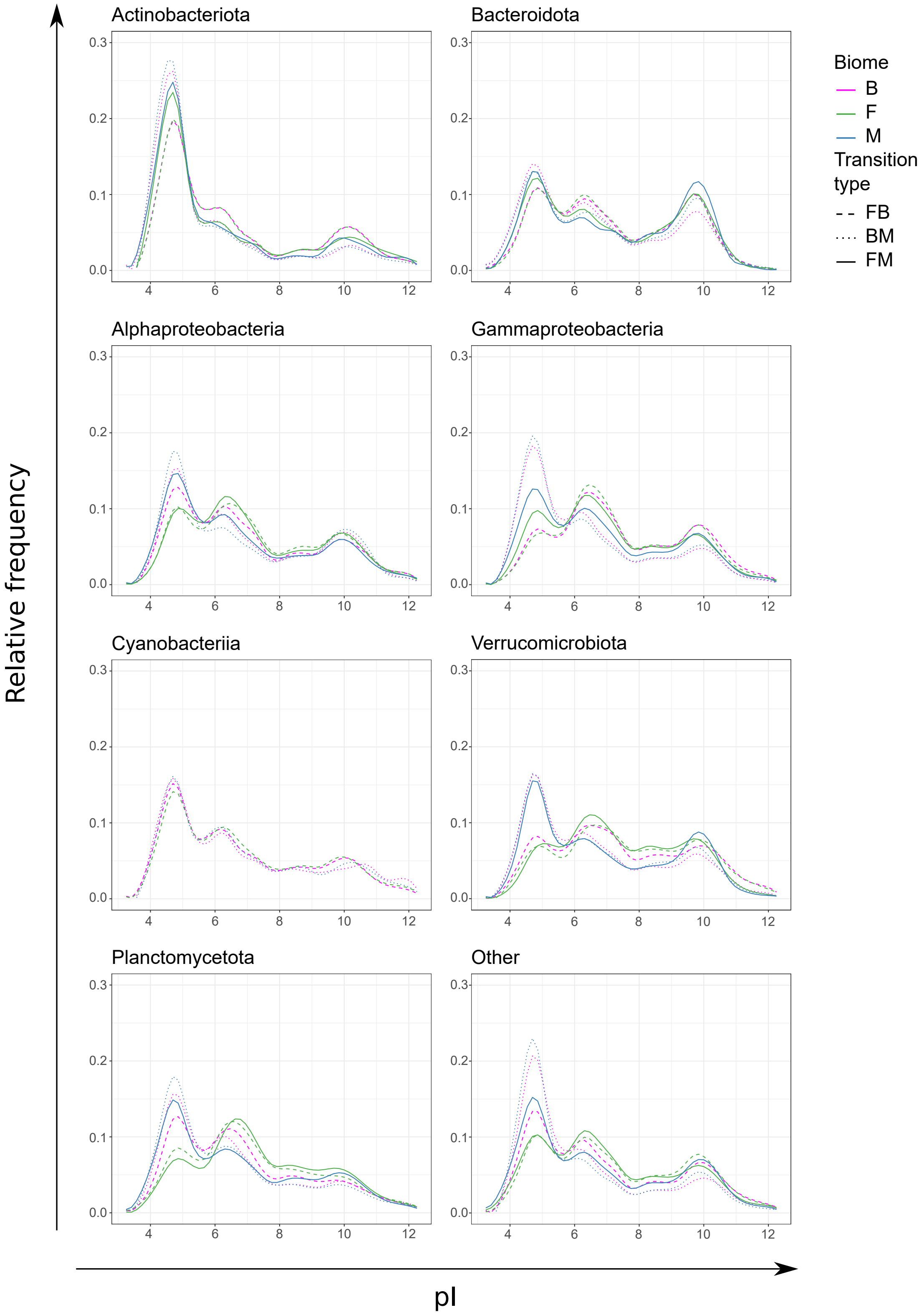


**Supplementary Fig. S5 | Averaged distribution of isoelectric point (pI) values in predicted proteomes of MAGs.** The distributions were averaged over all the MSGs from the same biome, transition type, and taxonomic group. The mean abundances of proteins within 0.5 pH wide bins (from pI ∈ [3.0, 3.5) to pI ∈ [12.0, 12.5)) were used to make the figure.

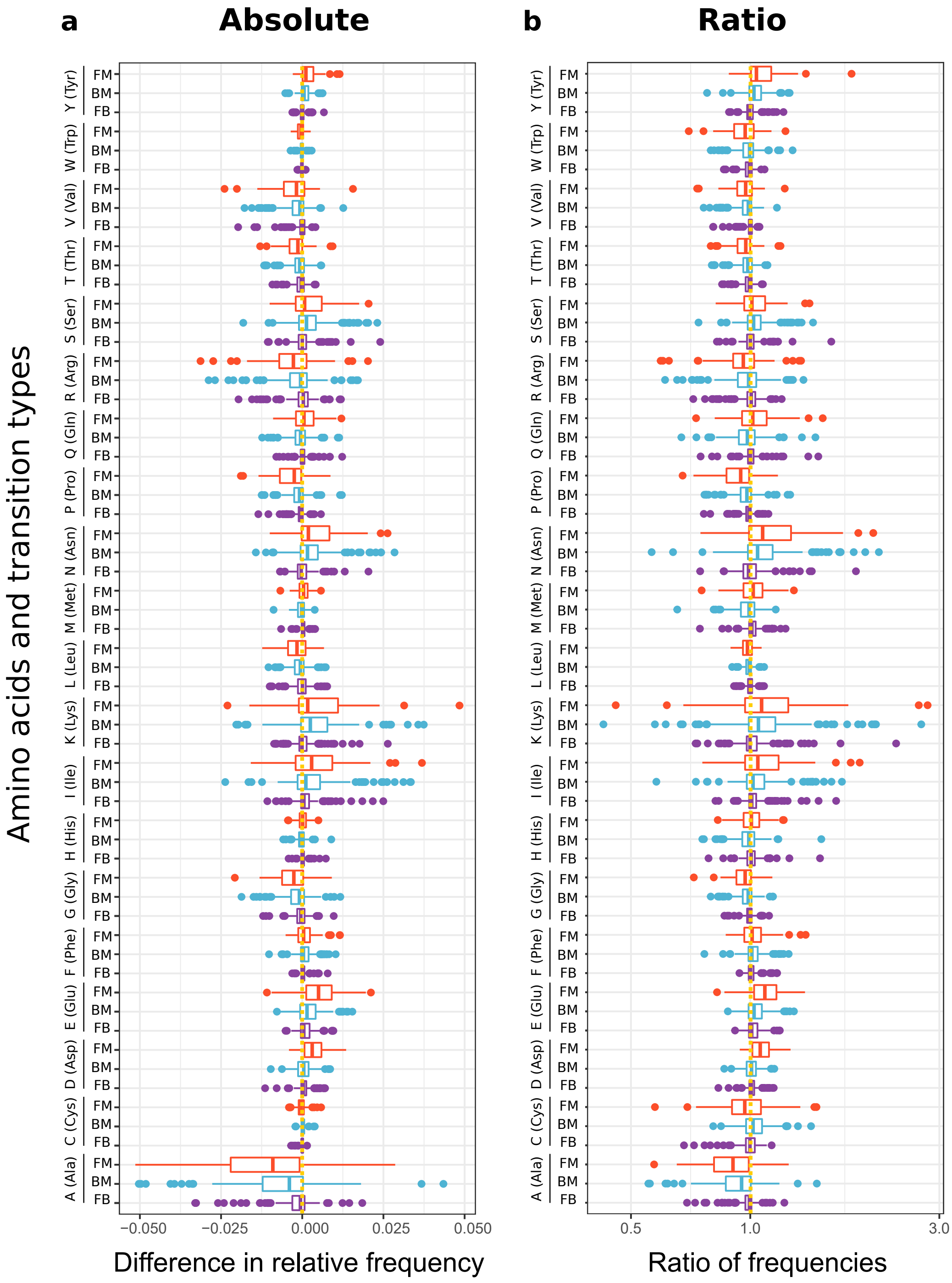


**Supplementary Fig. S6 | Differences in frequencies of amino acids in the proteomes across the transitions.** Presented as **a.** absolute differences in the relative frequencies or **b.** ratio of frequencies (logarithmic scale on x-axis).

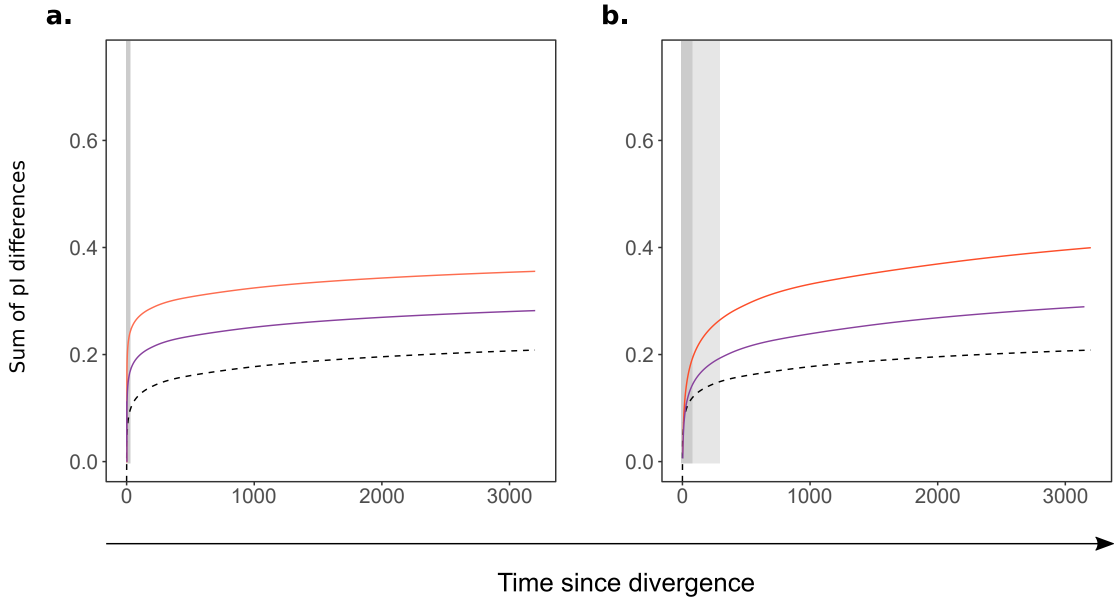


**Supplementary Fig. S7 | Models of proteome reorganization following cross-biome transitions.** The curves illustrate hypothetical dynamics of change in pI distribution after divergence events (as in Fig. 5d) connected to cross-biome transitions (coloured lines) or within the same biome (dashed black line). **a.** Abrupt change in the distribution of pIs driven by biome-specific selection occurs in the early stage after transition, followed by random changes occurring in the diverging lineages regardless of selective pressure (model A) **b.** The transition alters the rate at which the distributions of pIs change, which is observable over a long time span as a combined effect of random changes and selective pressure (model B). Fig. 5d does not allow to correctly distinguish between the models, as taxonomic biases and differences in distributions of times since divergence affect the fitted logarithmic models. However, differences in pI distributions between transitions and no-transition divergence events within the same taxonomic groups (as in Fig. 5g-i) allows distinguishing which dynamics are more prevalent. Under model A, differences in pI distribution between transitions and divergence events occurring shortly after them should be significant, corresponding to abrupt changes after transitions (gray background in **a.**). Regardless of the size of change induced by the transition (orange vs purple line), it should be distinguishable from random effects after a similarly short time span. On the other hand, under model B, the increased rate of changes would lead to systematic differences being distinguishable from the noise only after a significant amount of time. The difference in time needed for differences in pI distributions to be distinguishable between transition and "no-transition" events would depend on how much the rate of change increases after transitions. Under a highly increased rate of change (orange line), only in the early stages (dark-gray background in **b.**), the differences are indistinguishable from the noise. If the rate increase is weaker, it takes substantially longer for the differences to reach the same level (light-gray background in **b.**). Results shown in Fig. 5g-i suggest that model B is the most common dynamic in the bacterial world.

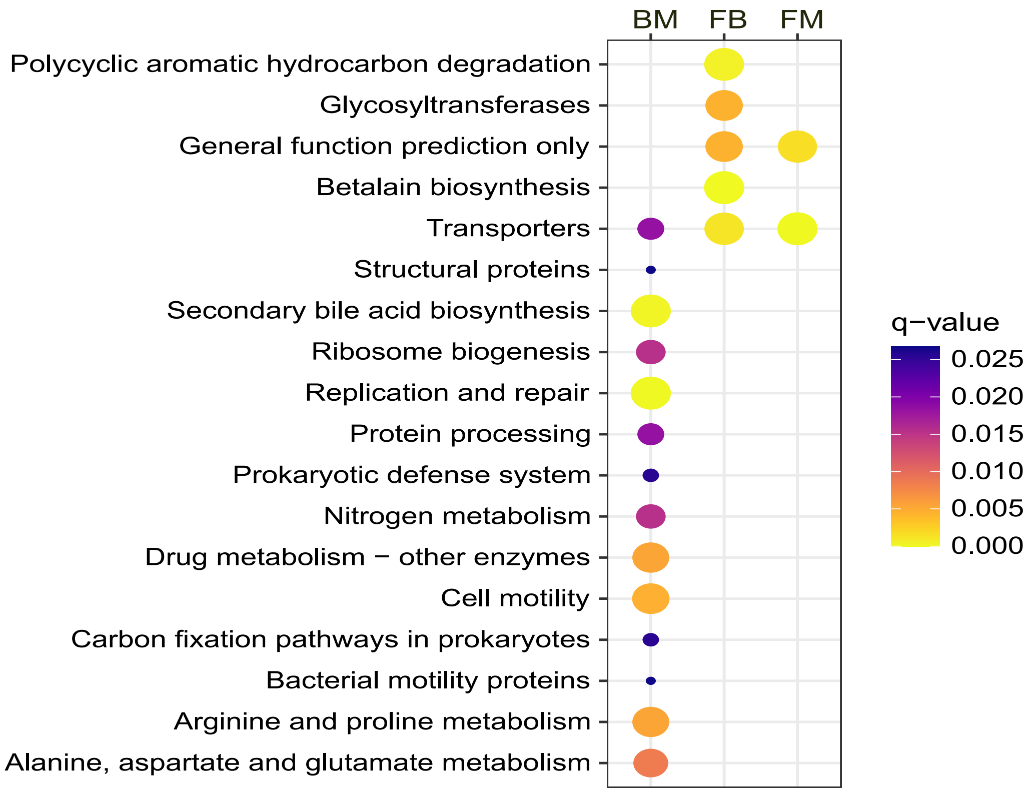


**Supplementary Fig. S8 | Results of over-representation analysis of the functional categories (KEGG Orthology level C) of differentially present KOs (i.e. gained/lost genes).** The probability of obtaining the observed number of KOs annotated to a category among the differentially present KOs, as if they were drawn at random from all the KOs analyzed for a transition type, was assessed using a hypergeometric test.

### Supplementary Discussion

Please note that information on "Role" and "Regulation" is mostly based on literature, while "Potential mechanism" is more speculative. In some cases, the genes' roles in adaptation or response to salinity have been previously described, and the citations are provided accordingly. Names of genes found in both FB and FM transitions are underlined, and the information for these are repeated since the direction of change was the same in all cases (from lower to higher salinity). Only the literature names from the cited papers are given and may refer to either the gene itself or its product (protein).

Please note that all the remarks on function and regulation come from the model organisms in which the genes were studied and may vary across the bacterial tree of life.

The roles in uninvestigated processes and regulatory processes are usually not mentioned. The lack of information is implied, though we encourage you to dive deeper into the literature when needed. Overall, do not treat these notes as conclusions on why these genes are differentially present across MSGs but rather as hypotheses. We hope these notes can serve as a starting for anybody interested in further investigations into the potential role of these genes in adaptation to salinity and beyond.

Abbreviations: MAG – metagenome-assembled genome; MSG – monobiomic sister group; FB – freshwater ↔ brackish; BM – brackish ↔ marine; FM – freshwater ↔ marine

#### Freshwater ↔ Brackish

##### K01531: P-type $Mg^{2+}$ transporter

More often present in MAGs from **freshwater** MSGs.
**Literature names:** MgtA/B
**Role:** $Mg^{2+}$ transport down the electrochemical gradient despite being an ATP-ase (Maguire 2006). Two different proteins in *Salmonella typhimurium* with very similar functions but significant differences in sequences (Tao et al. 1995).
**Regulation:** Expression induced by low concentrations of $Mg^{2+}$ (Tao et al. 1995; Maguire 2006)
**Potential mechanism:** Magnesium is one of the major components of sea salt and balancing its transport is one of the major challenges for bacteria transitioning across the salinity gradient (Walsh, Lafontaine, and Grossart 2013). Transporters expressed in response to low concentrations of $Mg^{2+}$ are expected to be more needed in freshwater, where we more often find them in the MAGs. Much remains unknown about the biochemistry and physiological role of these transporters (Maguire 2006).

##### *K15777: 4,5-DOPA dioxygenase extradiol

More often present in MAGs from **freshwater** MSGs.
**Literature names:** YgiD

**Role:** "The formation of betalamic acid from the precursor amino acid 3,4-dihydroxy-l-phenylalanine (l-DOPA)" (Gandía-Herrero and García-Carmona 2014). Betalains are a group of pigments with strong antiradical activity which can display different absorption spectra (Gandía-Herrero, Escribano, and García-Carmona 2010).
The gene has been found in freshwater single-cell genomes but without the rest of the betalain synthesis pathway genes (Ghylin et al. 2014) Extradiol dioxygenases cleave aromatic rings using molecular oxygen (Lipscomb 2008). The enzyme can also cleave 2,3-extradiol bonds and be involved in different synthesis pathways (Mueller, Hinz, and Zryd 1997).
**Regulation:** Unknown in bacteria (Guerrero-Rubio, García-Carmona, and Gandía-Herrero 2020).
**Potential mechanism:** Freshwater contains more dissolved organic matter, which causes scatters blue light, shifting the spectrum to longer wavelengths (Hill et al. 2019) and there might be a need for alternative pigments related to transitions in bacteria.

#### *K04100: protocatechuate 4,5−dioxygenase, alpha chain

More often present in MAGs from **freshwater** MSGs.
**Literature names:** ligA
**Role:** Involved in protocatechuate 4,5-cleavage pathway (Noda et al. 1990), involved in degradation of various aromatic compounds, including lignin-derived compounds (Kamimura and Masai 2014).
**Regulation:** Expressed when glucose is absent AND respective aromatic compounds are present (Tsagogiannis et al. 2021).
**Potential mechanism:** Allowing use of alternative sources of energy and carbon from dissolved organic matter.
**Note on co-annotation:** K04100 was the higher-confidence annotation in all cases. There were also genes annotated to K04100 and not to K15777, but not the other way around. Thus, we suggest putting more significance on K04100. However, the genes are related to each other (Burroughs et al. 2019) and both potential mechanisms relate to dissolved organic matter levels. Focused phylogenetic studies of the genes in specific MAGs could help to guide further investigations into actual mechanism at play.

#### K03498: trk/ktr system potassium uptake protein

More often present in MAGs from **brackish** MSGs.
**Literature names:** ktrB/D, trkH/G
**Role:** Part of a potassium uptake system crucial for responses to osmotic stress (Holtmann et al. 2003; Berry et al. 2003; Nanatani et al. 2015). When two system are present, the transporter is part of the one with lower affinity (Holtmann et al. 2003). Transports also $Rb^+$.
**Regulation:** Constitutive transcription (Holtmann et al. 2003), protein activity inhibited by c-di-AMP (Gibhardt et al. 2019).
**Potential mechanism:** Connected before with adaptation to higher salinity, allows long term growth in environment with higher osmolarity, but does not play role in responses sudden increase in osmotic pressure (Holtmann et al. 2003)

#### K03499: trk/ktr system potassium uptake protein

More often present in MAGs from **brackish** MSGs.
**Literature names:** ktrAC, trkA
**Role:** Part of a potassium uptake system crucial for responses to osmotic stress (Holtmann et al. 2003; Berry et al. 2003; Nanatani et al. 2015). When two system are present, the transporter is part of the one with higher affinity (Holtmann et al. 2003). Traports also $Rb^{+}$.
**Regulation:** Constitutive transcription (Holtmann et al. 2003), protein activity inhibited by c-di-AMP (Bai et al. 2014).
**Potential mechanism:** Connected before with adaptation to higher salinity, allows both survival of sudden increase in osmotic pressure as well as long term growth in environment with higher osmolarity (Holtmann et al. 2003).

#### K07301: Na+/Ca2+ antiporter

More often present in MAGs from **brackish** MSGs.
**Literature names:** YrbG
**Role:** $Na^{+}/Ca^{2+}$ antiporter, $Na^{+}$-coupled transport of $Ca^{2+}$ into membrane vesicles (Besserer et al. 2012)
**Regulation:** In *Escherichia coli* it is located on an operon with genes involved in outer membrane biogenesis. The operon is regulated by $\sigma^{E}$-dependent LPS stress signaling pathway (Martorana et al. 2011).
**Potential mechanism:** The $Na^{+}$-coupled transport conveyed by the antiporter is not energetically feasible in freshwaters because of the low extracellular $Na^{+}$ levels.

#### K16055: trehalose 6−phosphate synthase/phosphatase

**Literature names:** OtsB
**Role:** "an essential enzyme in the trehalose biosynthesis OtsAB pathway, catalyzes the dephosphorylation of trehalose-6-phosphate (trehalose-6-P) to generate trehalose, and plays a critical role in M. tuberculosis survival-associated cell wall formation and permeability" (Shan et al. 2016). Trehalose is a well-known osmoprotectant, accumulated inside the cell under osmotic stress by diverse bacteria and beyond (Ruhal, Kataria, and Choudhury 2013). It is also a component of cell-wall glycolipids, where it contributing to formation of a protective coat, as well as involved in transport of mycolic acids, which make cell wall less permeable (Shan et al. 2016). Allows bacteria to manage environmental stresses, including osmotic, thermal and oxidative ones (Hubloher et al. 2020).
**Regulation:** Multiple environmental stresses, including high salinity and temperatures, induce expression of the gene (Zeidler et al. 2017; Hubloher et al. 2020).
**Potential mechanism:** Has been shown to allow growth of *Escherichia coli* in increased salt concentrations, and of *Acinetobacter baumannii* when both salinity and temperature increase (Zeidler et al. 2017). Interestingly, in *Escherichia coli* the use of trehalose seems to depend on overproduction of trehalose and its degradation by periplasmatic trehalase, with the resulting glucose than being reutilized (Strøm and Kaasen 1993). Trehalases have been shown to be more often present in marine *Flavobacteriaceae* than in their marine relatives (Zhang et al. 2019). Potentially promotes adaptive phenotypic plasticity allowing life in different osmolarity/salinity.

Decreased cell-wall permeability may also explain why we do not see the same traces of brackish MGEs (mobile genetic elements) for FB transitions as for BM.

##### <u>K08974: putative membrane protein</u>

More often present in MAGs from **brackish** MSGs.
**Literature names:**
**Role:** Unknown. An analysis using STRING (Szklarczyk et al. 2019) to MEP (non-mevalonate) isoprenoid synthesis. STRING textmining conext it with only one article, which is about synthesis of isoprenoid pigments (Heider et al. 2014).
**Regulation:** Unknown.
**Potential mechanism:** Possibly related to changes in pigmentation due to shift of light spectra towards longer wavelength (red light) in less saline water (see K15777 for more details).

#### Brackish ↔ Marine

##### K01480: agmatinase

More often present in MAGs from **brackish** MSGs.
**Literature names:** SpeB (agmatine ureohydrolase)
**Role:** Polyamine biosynthesis. Catalyses reaction: agmatine + $H_2O$ ⇌ putrescine + urea. The reaction is part of one of the three polamine synthesis pathways, this one being utilized mostly by bacteria (Chitrakar et al. 2021). Polyamines are a group of compounds produced by organisms from all major evolutionary lineages, with diverse, complex and to large extent unexplained functions (Miller-Fleming et al. 2015). Results from cyanobacteria suggest key role of the enzyme in nitrogen metabolism (Burnat and Flores 2014). *Mycolicibacterium smegmatis* has the enzyme but does not produce putrescine (Zamakhaev et al. 2020), suggesting other functions of the enzyme or specific regulation of its activity. In plants, the polyamines allow to accommodate to quick changes in osmotic pressure (Kotakis et al. 2014).
**Regulation:** Induced by agmatine, repressed by cAMP (Szumanski and Boyle 1992).
**Potential mechanism:** Controlled production of polyamines can be a mechanism of plastic response to changes in salinity, characteristic to brackish waters as opposed to marine environment with relatively stable salt concentrations. In some cases, may be connected to post-translational modifications through K00809. Potentially promotes adaptive phenotypic plasticity allowing life in different osmolarity/salinity.
Use of urea as nitrogen source has been observed to be gained in specific groups of bacteria transitioning to more N-limited environments with lower salinity (Ramachandran, McLatchie, and Walsh 2021; Lanclos et al. 2022). Thus, the polyamine-producing pathway may also be used in opposite than conventional direction under N deficiency. Reactions and identified genes responsible for them:
(1) putrescine + urea ⇌ agmatine + $H_2O$ (K01480)
(2) agmatine + $CO_2$ ⇌ L-arginine (K01585 + potential role of carbonic anhydrase (K01673) to concentrate $CO_2$)

#### K04759: ferrous iron transport protein B

More often present in MAGs from **brackish** MSGs.
**Literature names:** FeoB
**Role:** The transporter part of the Feo $Fe^{2+}$ uptake system (Kammler, Schön, and Hantke 1993).
**Regulation:** Cytoplasmatic GTPase domain is believed to regulate transport (Marlovits et al. 2002). The *in vivo* active form is probably the complex with FeoA and FeoC (Stevenson, Wyckoff, and Payne 2016). All genes of Feo system in one operon believed to be regulated by metal availability (Cartron et al. 2006).
**Potential mechanism:** While Feo system is present in many marine genomes, it is often present exclusively with other iron uptake systems, especially $Fe^{3+}$ transporters (Hopkinson and Barbeau 2012). FeoB structure has been shown to be especially sensitive to excess salinity (Sestok, O'Sullivan, and Smith 2022). Change in salinity can also inhibit Fe-oxidation (or decrease in it can enable acquisition of brackish/freshwater oxidation pathways) (Cameron, Jones, and Edwards 1984). It may be that bacteria switch between different iron transporters due to these pressures.
A confounding factor might be the extent of hypoxia in the Baltic Sea, and samples from bigger depth from this basin, meaning the bacteria in question might have access to more $Fe^{2+}$, which is scarce in surface marine environments.

#### K00809: deoxyhypusine synthase

More often present in MAGs from **brackish** MSGs.
**Literature names:**
**Role:** Identified in eukaryotes and Archaea, where it is responsible for hypusination of IF-5A. However, bacteria do not have IF-5A. The traces of horizontal gene transfer of deoxyhypusine synthases from Archeae to bacteria has long known (Brochier, López-García, and Moreira 2004), yet the role of the transferred genes remains unexplained. A second substrate, outside of the protein, in hypusination is spermidine, a polyamine synthesis of which requires K01480 and K01585.
**Regulation:**
**Potential mechanism:** Possibly connects polyamine synthesis to post-translational modifications. Can influence charges on protein surface.

#### K01585: arginine decarboxylase

More often present in MAGs from **brackish** MSGs.
**Literature names:** SpeA
**Role:** Decarboxylases arginine to agmatine. Thus, the enzyme is directly upstream to K01480 in polyamine synthesis (Chitrakar et al. 2021) (see K01480 for more on polyamines).
**Regulation:** Inhibited by cAMP, repressed by putrescine (downstream product of polyamine synthesis) (Moore and Boyle 1991).
**Potential mechanism:** Controlled production of polyamines can be a mechanism of plastic response to changes in salinity, characteristic to brackish waters as opposed to marine

environment with relatively stable salt concentrations. May be connected to post-translational modifications through K00809.
Use of urea as nitrogen source has been observed to be gained in specific groups of bacteria transitioning to more N-limited environments with lower salinity (Ramachandran, McLatchie, and Walsh 2021; Lanclos et al. 2022). Thus, the polyamine-producing pathway may also be used in opposite than conventional direction under N deficiency. Reactions and identified genes responsible for them:
(1) putrescine + urea ⇌ agmatine + $H_2O$ (K01480)
(2) agmatine + $CO_2$ ⇌ L-arginine (K01585 + potential role of carbonic anhydrase (K01673) to concentrate $CO_2$)

#### K03782: catalase−peroxidase

More often present in MAGs from **brackish** MSGs.
**Literature names:** katG
**Role:** Can act as catalase – break up $H_2O_2$ to water and oxygen – as well as a peroxidase, an enzyme which uses $H_2O_2$ to oxidize other compounds. The latter function is crucial for degradation of many xenobiotics, and thus abundance of this gene in metagenomes has been connected to levels of water contamination (Muñoz-García et al. 2019), especially by polycyclic aromatic hydrocarbons (Witt 1995). Involved in antibiotic resistance against isoniazid (Muñoz-García et al. 2019).
**Regulation:** Expression inducible by stresses inducing production of reactive oxygen, such as heat shock (Godočíková et al. 2010).
**Potential mechanism:** Brackish enclosed basins accumulate higher concentrations of many xenobiotic substances than open oceans and the Baltic Sea has been in recent times reported to have up to 10 times higher concentrations of polycyclic aromatic hydrocarbons than the Northern Sea (Witt 1995). Thus, in these brackish environments recent pressures on acquisition of genes, rather than transition-related changes, might be the reason behind the difference in gene presence between the MSG pairs.
Could also be involved degradation of organic compounds coming from dissolved organic matter, or produced throughout their degradation. These are generally more diverse and present in higher concentrations in brackish water due to terrestrial input.

#### K21498: antitoxin HigA−1

More often present in MAGs from **brackish** MSGs.
**Literature names:** higA-1
**Role:** Part of toxin-antitoxin system stabilizing a superintegron in bacterial genomes (Christensen-Dalsgaard and Gerdes 2006).
**Regulation:** Expression induced by amino acid starvation (Christensen-Dalsgaard and Gerdes 2006).
**Potential mechanism:** Sign of difference in the mobile genetic elements which have spread and/or are being spread in the biomes: the superintegron seems to be more common in the brackish than marine environments. Genome streamlining in oligotrophic marine environments may also play role.

#### K19159: antitoxin YefM

More often present in MAGs from **brackish** MSGs.
**Literature names:** yefM
**Role:** Part of toxin-antitoxin system stabilizing yefM-yoeB cassette in bacterial genomes (Kędzierska, Lian, and Hayes 2007).
**Regulation:** Autorepression (Kędzierska, Lian, and Hayes 2007).
**Potential mechanism:** Sign of difference in the mobile genetic elements which have spread and/or are being spread in the biomes: the cassette seems to be more common in the brackish than marine environments. Genome streamlining in oligotrophic marine environments may also play role.

#### K07064: uncharacterized protein

More often present in MAGs from **brackish** MSGs.
**Literature names:**
**Role:**
**Regulation:**
**Potential mechanism:** Uncharacterized protein. Its clustering with the antitoxins could guide characterization.

#### #K05844: ribosomal protein S6−−L−glutamate ligase

More often present in MAGs from **brackish** MSGs.
**Literature names:** rimK
**Role:** Modification of ribosomal protein S6 and synthesis of poly- α-glutamic acid (Kino, Arai, and Arimura 2011). Involved in regulation of motility, surface attachment and rhizosphere colonization (Little et al. 2016).
**Regulation:** Complex regulatory system involving cyclic-di-GMP, expression induced by cold and starvation (salinity not assessed) (Grenga et al. 2020).
**Potential mechanism:** Less specific protein modifications or regulated production of intracellular osmolytes as plastic responses to changing salinity levels.
**Note on co-annotation:** K05844 was the highest-confidence annotation in all cases. There were also genes annotated to K05844 and not to K18310/ K14940, but not the other way around. Thus, we suggest putting more significance on K04100. However, the genes are homologs (Li et al. 2003; Collard et al. 2010) and the actual genes might well be other bacterial paralogs of K04100. Focused phylogenetic studies of the genes in specific MAGs could help to guide further investigations into actual mechanism at play.

#### #K18310: beta−citrylglutamate/N−acetylaspartylglutamate synthase

More often present in MAGs from **brackish** MSGs.
**Literature names:** RIMKLB
**Role:** In mammals catalases synthesis of N-acetylaspartylglutamate and β-citrylglutamate (Collard et al. 2010).
**Regulation:**

**Potential mechanism:** Probably misanotation due to similarity with K05844 and K14940. Seems that KEGG has different annotation to bacterial, archeal and eukaryotic homologs, since they perform different functions and whether they are orthologs or paralogs might be difficult to distinguish. May suggest that actual gene functions are similar but not the same as for any of the genes, potentially including other pathways and protein modifications involving glutamic acid or glutamate.

#### #K14940: gamma−F420−2:alpha−L−glutamate ligase

More often present in MAGs from **brackish** MSGs.
**Literature names:** cofF
**Role:** In methanogenic archeon *Methanococcus jannaschii* it is a glutamate ligase modifying methanogenic coenzyme F420 (Li et al. 2003).
**Regulation:**
**Potential mechanism:** Possibly misanotation due to similarity with K05844, however horizontal transfer from *Archeae* cannot be excluded, if the gene can find alternative function in bacterial cells. Seems that KEGG has different annotation to bacterial, archeal and eukaryotic homologs, since they perform different functions and whether they are orthologs or paralogs might be difficult to distinguish. May suggest that actual gene functions are similar but not the same as for any of the genes, supporting hypothesis that modifications of other targets than ribosomal protein S6 can be at play.

#### K03284: magnesium transporter

More often present in MAGs from **brackish** MSGs.
**Literature names:** corA
**Role:** Passive transport of $Mg^{2+}$, as well as other ions such as $Co^{2+}$, $Ni^{2+}$ and $Zn^{2+}$ (Stetsenko and Guskov 2020). One of the major and most common bacterial magnesium uptake systems, and opposed to others, can facilitate also $Mg^{2+}$ efflux (Maguire 2006).
**Regulation:** Gated transport regulated by intracellular $Mg^{2+}$ levels (Payandeh, Pfoh, and Pai 2013).
**Potential mechanism:** Magnesium is a major component of the sea salt and decreasing salinity may deem need for acquisition for alternative/additional magnesium uptake systems. Clustering with K03282 suggests that the transporter may play a role in either managing hypoosmotic stress (through ion efflux) or recovery from it (by replenishing the pool of $Mg^{2+}$ and other transported cations). Thus, potentially allows adaptive phenotypic plasticity allowing life in different osmolarity/salinity.

#### K03282: large conductance mechanosensitive channel

More often present in MAGs from **brackish** MSGs.
**Literature names:** mscL
**Role:** Mechanosensitive channel involved in responses to hyperosomotic shock, preventing turgor pressure which would destroy the cell, allowing accommodation to new conditions and further growth (Levina et al. 1999). It has the highest conductance among *Escherichia coli* mechanosensitive channels (Levina et al. 1999), thus is responsible for the strongest response to hypoosomotic stress.

**Regulation:** Mechanosensitive channel, activity regulated by tension on the membrane.
**Potential mechanism:** Directly connected to hypoosmotic stress, possible experience in the course of marine to brackish transitions. Brackish bacteria experience bigger changes in salinity of their immediate environment. Potentially allows adaptive phenotypic plasticity allowing life in different osmolarity/salinity.

#### K02237: competence protein ComEA

More often present in MAGs from **brackish** MSGs.
**Literature names:** comEA
**Role:** DNA receptor and involved in DNA uptake in naturally competent bacteria (Seitz et al. 2014).
**Regulation:** Competence is initiated in response to various environmental cues, with the relevant cues and the response to them differing widely among bacteria (Seitz and Blokesch 2013). However, nutrient limitation and cell density may be listed among the most common ones (Seitz and Blokesch 2013).
**Potential mechanism:** Two components of the competence system (see also K02238), as well as a regulator of competence (K07343), all are differentially present across the BM MSG pairs. Another of the identified genes, K03630, was originally thought to take part competence but it was later shown to be disposable for the process, though its expression is induced in competent bacteria. Therefore, natural competence is probably more important for adaptation to brackish conditions than results for the single genes suggest. The additional role of K02238 in regulation type IV pili expression may explain a different pattern based on presence/absence changes than for K02237, which is also generally present in less of the MAGs.
Brackish environments are less stable than open oceans and pose pressure for adaptation to wider salinity spectrum. Brackish bacteria might benefit from higher genomic plasticity, as it allows adaptation to disruptions in the ecosystem and colonization of different niches across the salinity gradient (through acquisition of genes shifting optimal/tolerated salt concentrations).
Anthropogenic pollution may be a confounder, as horizontal gene transfer has probably been crucial mechanism of adaptation to emergence of xenobiotics in the environment (Springael and Top 2004).

#### K06987: uncharacterized protein

More often present in MAGs from **brackish** MSGs.
**Literature names:**
**Role:**
**Regulation:**
**Potential mechanism:** Uncharacterized protein. Its clustering with the competence protein ComEA (K02237) could guide characterization.

#### K03321: sulfate permease, SulP family

More often present in MAGs from **brackish** MSGs.
**Literature names:** sulP
**Role:** Diverse group of transporters, mainly anion:cation symporters and anion:anion antiporters, often fused to other proteins, including carbonic anhydrases (Felce and Saier

2004). Includes characterized $Na^+$-coupled $HCO_3^-$ transporters (Price et al. 2004), as well as $Na^+$-independent sulfate transporters (Zolotarev et al. 2008), which may play role in sulfate transport for antibiotic production (Alper and Sharma 2013).
**Regulation:** Probably varies widely in different organisms and depending on the actual role of the ortholog.
**Potential mechanism:** Even though $Na^+$-coupled, $HCO_3^-$ transport may be important function for adaptation, especially as we also have a carbonic anhydrase (K01673) preferentially present in brackish MAGs across BM MSG pairs. It might be a matter of affinity or regulation that additional hydrocarbonate uptake system is needed in brackish bacteria. Potentially its interactions with carbonic anhydrase play a role.
Sulfate transport for antibiotic production may be of importance as bacteria move into more eutrophic brackish waters. Especially as the Baltic can be now expected to be inhabited by more species adapted to living in eutrophic conditions, drastic forms of which have been induced by human activity in the last decades.

#### K02036- K02040 phosphate transport system

More often present in MAGs from **brackish** MSGs.
Described together as they act together forming a complex, and we observed them to be almost always gained/lost/present together.
**Literature names:** pst
**Role:** ATP-dependent phosphate uptake system (Webb, Rosenberg, and Cox 1992), crucial for responses to phosphate deficiency. In some organisms it can be a sensor for transcriptional regulation of genes related to phosphate deficiency (Suzuki et al. 2004; Västermark and Saier 2014), as well as for formation of survival cell forms (Namugenyi et al. 2017)
**Regulation:** Expression is induced in when phosphate availability is low (Qi, Kobayashi, and Hulett 1997). $Na^+$ can also induce its transcription, while availability of energy sources and pH can regulate activity of the protein (Burut-Archanai, Eaton-Rye, and Incharoensakdi 2011)
**Potential mechanism:** Marine bacteria often use $Na^+$-coupled phosphate transport (Walsh, Lafontaine, and Grossart 2013), which becomes infeasible with decreasing salinity. Responses to environmental cues and ability to stimulate formation of survival forms may play additional role in allowing phenotypic plasticity in response to unstable brackish conditions and during transitions.

#### K01673: carbonic anhydrase

More often present in MAGs from **brackish** MSGs.
**Literature names:** can, cynT
**Role:** Converts bicarbonate into $CO_2$ and the other way around (more precisely catalyses reaction $HCO_3^- + H^+ \leftrightarrow CO_2 + H_2O$). Carbonic anhydrases are used for both concentration and disposal of $CO_2$, as well as to provide sufficient concentrations of $HCO_3^-$/ $CO_2$ for enzymatic reactions (Smith and Ferry 2000). In favorable conditions $HCO_3^-$ transport is primally conveyed through a specialized transporter (Fan et al. 2021), which is a $HCO_3^-/Na^+$ symporter (Fan et al. 2019).
**Regulation:** Increased expression in response to stresses: heat, starvation, high cell density (Merlin et al. 2003).
**Potential mechanism:** In animals, anhydrase activity is induced by and is crucial for accommodation to lower salinity, as it compensates for decrease in $CO_2$ disposal through

$HCO_3^-/Cl^-$ antiporters (Henry 2001). In bacteria, $Na^+$ coupled $HCO_3^-$ import seems to be more important (Fan et al. 2019; 2021), but the situation is analogous. It may be that marine, but not brackish bacteria can rely solely on $Na^+$ coupled transport. Role of anhydrase in accommodation to various stresses may also be important in variable and less stable brackish environment. Potentially promotes adaptive phenotypic plasticity allowing life across environmental gradients in the brackish biome.
Use of urea as nitrogen source has been observed to be gained in specific groups of bacteria transitioning to more N-limited environments with lower salinity (Ramachandran, McLatchie, and Walsh 2021; Lanclos et al. 2022). Thus, the polyamine-producing pathway may also be used in opposite than conventional direction under N deficiency. Reactions and identified genes responsible for them:
(1) putrescine + urea ⇌ agmatine + $H_2O$ (K01480)
(2) agmatine + $CO_2$ ⇌ L-arginine (K01585 + potential role of carbonic anhydrase (K01673) to concentrate $CO_2$)

#### K07007: 3−dehydro−bile acid Delta4,6−reductase

More often present in MAGs from **brackish** MSGs.
**Literature names:** baiN
**Role:** Identified in a gut bacterium, hydrogenates double bonds between carbon atoms, most notably in isoprenoids and steroids (Harris et al. 2018). In gut, it is believed to convey a crucial reductive step of secondary bile acid synthesis by bacteria in the large bowel (Harris et al. 2018). The role in free-living bacteria unknown.
**Regulation:** Not on the same operon with other secondary bile acid synthesis genes, unknown mechanisms of regulation (Doden and Ridlon 2021).
**Potential mechanism:** As in the case of K15777/K04100 and K08974 for FB (and FM in the latter case) transitions, the difference in dissolved organic carbon sources may be related to differential presence of this gene. As the enzyme can modify isoprenoids and steroids, it can play role in synthesis of alternative pigments connected to shift in light spectrum towards longer wavelengths. It may also have uncharacterized roles in metabolism of alternative carbon sources and/or xenobiotic degradation, the latter possibly connected with pollution in the enclosed brackish basins.

#### K07290: AsmA family protein

More often present in MAGs from **brackish** MSGs.
**Literature names:** yhjG
**Role:** Related asmA proteins have a role **"**preventing misfolding of outer membrane proteins" (Prieto et al. 2009; based on: Misra and Miao 1995; Deng and Misra 1996). It is also a homolog of eukaryotic lipid transfer proteins (Levine 2019)
**Regulation:**
**Potential mechanism:** If this protein is involved in control of misfolding of outer membrane proteins, this process might be crucial as bacteria move across salinities in the brackish biome, which would change the folding dynamics of proteins exposed to external environment.
**Note:** The gain/loss of this gene, unlike almost all others, is taxonomically confined to *Planctomycetota* and *Proteobacteria*.

#### K07115: 23S rRNA (adenine2030−N6)−methyltransferase

More often present in MAGs from **brackish** MSGs.
**Literature names:** rlmJ
**Role:** Catalyses specific modification of adenine 2030A in 23S rRNA to methyloadenine (Golovina et al. 2012). It has been suggested that "N6 methylation of adenosine may enhance long-range stacking interactions" (Sergiev et al. 2016; based on Kierzek and Kierzek 2003).
**Regulation:** Probably constitutive expression.
**Potential mechanism:** Change in salinity can have major effects on biochemistry and biophysics of biomolecules, as the proteome-scale changes in protein properties and amino acid composition suggest. Thus, the long-range stacking interactions may also be affected by salinity, even though they occur in relatively less affected intracellular environment. Marine genome streamlining may also be of importance.
**Note:** The gain/loss of this gene, unlike almost all others, is taxonomically confined to *Proteobacteria*.

#### K02482: NtrC family, sensor kinase

More often present in MAGs from **brackish** MSGs.
**Literature names:** flgS
**Role:** Part of FlgS/FlgR two-component signal transduction system, which regulates transcription of the fla regulon in *Campylobacter jejuni* (Wösten, Wagenaar, and van Putten 2004). The fla regulon contains flagellar biosynthesis proteins and its activation is associated with growth and motility (Wösten, Wagenaar, and van Putten 2004). In *Helicobacter pylori*, it responses to changes in pH (Wen et al. 2009).
**Regulation:** A sensor, changes in activity in response to environmental cues convey its function (see "Role:" section)
**Potential mechanism:** Induction of cell motility in conditions allowing for growth may be important for transitions, as well as moving along the gradients of the brackish biome and colonizing new/wider niches within it. Potentially promotes adaptive phenotypic plasticity allowing life across environmental gradients in the brackish biome. More information about environmental cues to which the sensor can respond is needed to hypothesize about its importance.

#### K07343: DNA transformation protein and related proteins

More often present in MAGs from **brackish** MSGs.
**Literature names:** tfox
**Role:** Together with another gene, HapR, it regulates expression of comEA (K02237), and thus the natural competence (Metzger et al. 2019). It also regulates expression of type VI secretion system and interbacterial killing (Metzger et al. 2019). It is activated in response to chitin, and it has been suggested to be crucial for colonization of chitinous surfaces by *Vibrio* species, at the same time reducing motility through repression of related protein TfoY (Metzger et al. 2019).
**Regulation:** Activated in response to chitin (Metzger et al. 2019).
**Potential mechanism:** Two components of the competence system (see also K02237 and K02238), together with this regulator of competence, all are differentially present across the BM MSG pairs. Another of the identified genes, K03630, was originally thought to take part

competence but it was later shown to be disposable for the process, though its expression is induced in competent bacteria. Therefore, natural competence is probably more important for adaptation to brackish conditions than results for the single genes suggest. The additional role of K02238 in regulation type IV pili expression may explain a different pattern based on presence/absence changes than for K02237, which is also generally present in less of the MAGs.

Brackish environments are less stable than open oceans and pose pressure for adaptation to wider salinity spectrum. Brackish bacteria might benefit from higher genomic plasticity, as it allows adaptation to disruptions in the ecosystem and colonization of different niches across the salinity gradient (through acquisition of genes shifting optimal/tolerated salt concentrations).

The importance of this gene for colonization of chitinous surfaces may suggest that associating to an animal host may have a role in transitions. Potentially, as animals adapt to a shift in salinity related to formation of a brackish basin, but local bacteria get outcompeted by the global brackish microbiome, association to a host is a survival strategy for the local bacteria. The host-associated bacteria are less likely to come from the other, distant brackish basins, and niches on the host surface open-up.

#### K03630: DNA repair protein RadC

More often present in MAGs from **brackish** MSGs.
**Literature names:** radC
**Role:** Originally connected to competence and UV-light protection (DNA repair), this connection has been put into question by results on *Streptococcus pneumoniae* (Attaiech et al. 2008). However, it might be that the gene present in *Streptococcus pneumoniae*, especially the UV-protecting properties, which have been shown in *Rhodobacter capsulatus* (Katsiou et al. 1999), ecologically more suspectable to UV-light.
**Regulation:** It is induced in competent bacteria (Attaiech et al. 2008) and in
**Potential mechanism:** Might be connected to repair of transfer DNA in competent bacteria, though this role has been put into question and there is evidence against it, though limited to *Streptococcus pneumoniae* (Attaiech et al. 2008).

The role in DNA repair in response to UV light is opposite to the light wavelength shift observed in relation to higher dissolved organic carbon levels in lower salinities, as the organic particles scatter short-length light waves. However, clustering with genes responsible for chemotaxis gives a possible explanation. As bacteria move, lead by environmental cues, they may migrate to different higher parts of the water column, exposing themselves to stronger irradiation.

#### K03413: chemotaxis family, chemotaxis protein CheY

More often present in MAGs from **brackish** MSGs.
**Literature names:** CheY
**Role:** "Diffusible response regulator", key gene for chemotaxis based on flagellar motion (Colin et al. 2021). CheY proteins can respond to multiple and various environmental cues, even within one bacterial cell (Ganusova et al. 2021).
**Regulation:** Activity regulated in response to environmental cues.
**Potential mechanism:** Gain/loss of this protein strongly points towards changes in chemotaxis. The diversity of potential cues to which it responds make it hard to speculate

about the specific mechanism. However, it is highly probable cheY genes responding to different factors are gained by different bacteria, and its differential presence points towards importance of responses to the gradients of environmental factors, which are characteristic of most brackish environments and usually much more pronounced there than in the open ocean. Potentially promotes adaptive phenotypic plasticity allowing movement across environmental gradients in the brackish biome.
The fact that this gene clusters with weakly characterized sensor histidine kinase (K20974) involved in motility, instead of the literature partner cheA (K03407, not among identified genes), suggests alternative signal transduction pathway to be at play.

#### K20974: two−component system, sensor histidine kinase

More often present in MAGs from **brackish** MSGs.
**Literature names:** Hpt
**Role:** Histidine kinase, needed for swarming activity and biofilm formation (Hsu et al. 2008)
**Regulation:** As a hisitidine kinase it is involved in transducing signals in the cell, and its activity depends on the proteins with which it interacts.
**Potential mechanism:** This gene, involved in motility, clusters with CheY (K03413(, key chemotaxis protein. It suggests an alternative chemotaxis transduction pathway to be at play, using this kinase instead of usually described cheY partner, that is instead of the literature partner cheA (K03407, not among identified genes). It probably allows bacteria to respond to at least one of the environmental factor gradients, characteristic for brackish environments. Potentially promotes adaptive phenotypic plasticity allowing life and/or movement across environmental gradients in the brackish biome.

#### K07497: putative transposase and K07483: transposase

More often present in MAGs from **brackish** MSGs.
**Literature names:**
**Role:** Transposition, i.e. excision of a part of DNA molecule and insertion of it elsewhere (in the same or another molecule) .
**Regulation:**
**Potential mechanism:** Higher abundance of transposases in brackish (Baltic Sea) metagenomes, as compared to closely located marine environments, has been previously observed (Vigil-Stenman et al. 2017). Our results strengthen the notion of increased genomic plasticity in brackish bacteria.
Brackish environments are less stable than open oceans and pose pressure for adaptation to wider salinity spectrum. Brackish bacteria might benefit from higher genomic plasticity, as it allows adaptation to disruptions in the ecosystem and colonization of different niches across the salinity gradient (through acquisition of genes shifting optimal/tolerated salt concentrations).
**Note on coannotation:** This gene has been found in more cases than K07483. Have the latter (K07483) has been coannotated also as K07497 in less than 60% of the cases, suggesting that both are differential present across the MSG pairs regardless of the coannotation.

#### K02238: competence protein ComEC

More often present in MAGs from **brackish** MSGs.
**Literature names:** comEC
**Role:** Transports one strand of the imported DNA and degrades the other (Silale, Lea, and Berks 2021). It is also key for inducing expression of type IV pili (Salzer et al. 2016), which has a role in multiple aspects of bacterial physiology, including motility, cell adhesion and protein secretion (Melville and Craig 2013).
**Regulation:** Competence is initiated in response to various environmental cues, with the relevant cues and the response to them differing widely among bacteria (Seitz and Blokesch 2013). However, nutrient limitation and cell density may be listed among the most common ones (Seitz and Blokesch 2013).
**Potential mechanism:** Two components of the competence system (see also K02237 and K02238), as well as a regulator of competence (K07343), all are differentially present across the BM MSG pairs. Another of the identified genes, K03630, was originally thought to take part competence but it was later shown to be disposable for the process, though its expression is induced in competent bacteria.Therefore, natural competence is probably more important for adaptation to brackish conditions than results for the single genes suggest. The additional role of K02238 in regulation type IV pili expression may explain a different pattern based on presence/absence changes than for K02237, which is also generally present in less of the MAGs.
Brackish environments are less stable than open oceans and pose pressure for adaptation to wider salinity spectrum. Brackish bacteria might benefit from higher genomic plasticity, as it allows adaptation to disruptions in the ecosystem and colonization of different niches across the salinity gradient (through acquisition of genes shifting optimal/tolerated salt concentrations).

#### K07460: putative endonuclease

More often present in MAGs from **brackish** MSGs.
**Literature names:**
**Role:** Endonucleases are enzymes that cleave DNA into two different DNA molecules, as opposed to exonucleases, which cleave off nucleotides at the ends of DNA sequences. This protein to the best of our knowledge has not been characterized.
**Regulation:**
**Potential mechanism:** The relatively similar presence and gain/loss pattern to K02238 suggest possible role in competence. Connection to other mobile genetic elements, and thus genomic plasticity, is also possible.
Brackish environments are less stable than open oceans and pose pressure for adaptation to wider salinity spectrum. Brackish bacteria might benefit from higher genomic plasticity, as it allows adaptation to disruptions in the ecosystem and colonization of different niches across the salinity gradient (through acquisition of genes shifting optimal/tolerated salt concentrations).

#### K07391: magnesium chelatase family protein

More often present in MAGs from **brackish** MSGs.
**Literature names:** *yifB*

**Role:** A protease, probably with chaperon properties, i.e. degrading misfolded proteins (Iyer et al. 2004). Belongs to $Mg^{2+}$ chelatase family, though its potential function as a chelatase has not been, to the best of our knowledge, investigated.
**Regulation:** While little is known about the regulation of expression or activity of the protein, the gene is among ones frequently changed course of integration of mobile genetic elements into the genome, with consistent disruption of one domain (Mageeney et al. 2020). Considering that it clusters with possible mobile-genetic element related genes, it may be that it is the disrupted version of the gene that conveys the function.
**Potential mechanism:** As magnesium is a major component of sea salt, the chelatase activity could allow to stop the cations from leaving the cell. Chaperon function could be more beneficial after transitions and in changing salinity, as amino acid composition, mainly charges on the proteins, is then misadjusted to the new conditions, increasing the ratio of misfolded proteins.

### Freshwater ↔ Marine

#### K03549: KUP system potassium uptake protein

More often present in MAGs from **freshwater** MSGs.
**Literature names:** kup
**Role:** Potasium uptake system. Connected to hyperosmotic stress, has little impact on potassium transport at neutral pH but plays a crucial role in the acidic environment (Trchounian and Kobayashi 1999). $K^+/H^+$ symporter (Rodriguez-Navarro, Blatt, and Slayman 1986; Tascón et al. 2020)
**Regulation:** Activity depeneds on $K^+$ concentrations (Tascón et al. 2020).
**Potential mechanism:** Observed changes in the presence/absence of this gene may thus be related to the fact that pH is buffered at stable levels in the ocean and highly variable in freshwater. Freshwater bacteria experience osmotic pressure due to osmolates other than sea salt, in conditions which can often be connected with higher acidity. Might also represent a switch to $H^+$-coupled $K^+$ import, as opposed to ATP-dependent transport, which would be preferable and energetically feasible in freshwater. It might also be more feasible to use proton motive force directly, and not ATP, by freshwater but not marine bacteria, as for the latter the protons in the immediate surrounding would be buffered by the salts.

#### K08974: putative membrane protein

More often present in MAGs from **marine** MSGs.
**Literature names:**
**Role:** Unknown. An analysis using STRING (Szklarczyk et al. 2019) to MEP (non-mevalonate) isoprenoid synthesis. STRING textmining conext it with only one article, which is about synthesis of isoprenoid pigments (Heider et al. 2014).
**Regulation:** Unknown.
**Potential mechanism:** Possibly related to changes in pigmentation due to shift of light spectra towards longer wavelength (red light) in less saline water (see K15777 for more details).

#### K03499: trk/ktr system potassium uptake protein

More often present in MAGs from **marine** MSGs.
**Literature names:** ktrAC, trkA
**Role:** Part of a potassium uptake system crucial for responses to osmotic stress (Holtmann et al. 2003; Berry et al. 2003; Nanatani et al. 2015). When two system are present, the transporter is part of the one with higher affinity (Holtmann et al. 2003). Traports also $Rb^+$.
**Regulation:** Constitutive transcription (Holtmann et al. 2003), protein activity inhibited by c-di-AMP (Bai et al. 2014).
**Potential mechanism:** Connected before with adaptation to higher salinity, allows both survival of sudden increase in osmotic pressure as well as long term growth in environment with higher osmolarity (Holtmann et al. 2003).

#### K07301: Na+/Ca2+ antiporter

More often present in MAGs from **marine** MSGs.
**Literature names:** YrbG
**Role:** $Na^+/Ca^{2+}$ antiporter, $Na^+$-coupled transport of $Ca^{2+}$ into membrane vesicles (Besserer et al. 2012)
**Regulation:** In *Escherichia coli* it is located on an operon with genes involved in outer membrane biogenesis. The operon is regulated by $\sigma^E$-dependent LPS stress signaling pathway (Martorana et al. 2011).
**Potential mechanism:** The $Na^+$-coupled transport conveyed by the antiporter is not energetically feasible in freshwaters because of the low extracellular $Na^+$ levels.

#### K16052: MscS family membrane protein

More often present in MAGs from **marine** MSGs.
**Literature names:** ynaI
**Role:** Low-conductivity, $Na^+/K^+$ selective mechanosensitive channel (Yu et al. 2018).
**Regulation:** Mechanosensitive channel, activity regulated by tension on the membrane.
**Potential mechanism:** As it is a low-conductivity, $Na^+/K^+$ selective channel, it may be feasible to use it for regulated, low-cost management of hypoosmotic/turgor stress in marine, but not freshwater environments. Loss of $K^+$ may be faster to recover from in saline waters rich in potassium ions, though energetically costly (though potentially less than opening a channel with lower selectivity). Also, the use of sodium-motive force by marine bacteria (Walsh, Lafontaine, and Grossart 2013) may occasionally lead to uncontrolled accumulation of $Na^+$ inside the cell, which could be managed by the transporter.

Golovina, Anna Y., Margarita M. Dzama, Ilya A. Osterman, Petr V. Sergiev, Marina V. Serebryakova, Alexey A. Bogdanov, and Olga A. Dontsova. 2012. "The Last RRNA

Methyltransferase of E. Coli Revealed: The YhiR Gene Encodes Adenine-N6 Methyltransferase Specific for Modification of A2030 of 23S Ribosomal RNA." *RNA* 18 (9): 1725–34. https://doi.org/10.1261/rna.034207.112.
Grenga, Lucia, Richard Howard Little, Govind Chandra, Stuart Daniel Woodcock, Gerhard Saalbach, Richard James Morris, and Jacob George Malone. 2020. "Control of MRNA Translation by Dynamic Ribosome Modification." *PLoS Genetics* 16 (6): e1008837. https://doi.org/10.1371/journal.pgen.1008837.
Guerrero-Rubio, María Alejandra, Francisco García-Carmona, and Fernando Gandía-Herrero. 2020. "First Description of Betalains Biosynthesis in an Aquatic Organism: Characterization of 4,5-DOPA-extradiol-dioxygenase Activity in the Cyanobacteria Anabaena Cylindrica." *Microbial Biotechnology* 13 (6): 1948–59. https://doi.org/10.1111/1751-7915.13641.
Harris, Spencer C., Saravanan Devendran, João M. P. Alves, Sean M. Mythen, Phillip B. Hylemon, and Jason M. Ridlon. 2018. "Identification of a Gene Encoding a Flavoprotein Involved in Bile Acid Metabolism by the Human Gut Bacterium Clostridium Scindens ATCC 35704." *Biochimica et Biophysica Acta (BBA) - Molecular and Cell Biology of Lipids* 1863 (3): 276–83. https://doi.org/10.1016/j.bbalip.2017.12.001.
Heider, Sabine A. E., Natalie Wolf, Arne Hofemeier, Petra Peters-Wendisch, and Volker F. Wendisch. 2014. "Optimization of the IPP Precursor Supply for the Production of Lycopene, Decaprenoxanthin and Astaxanthin by Corynebacterium Glutamicum." *Frontiers in Bioengineering and Biotechnology* 2. https://www.frontiersin.org/article/10.3389/fbioe.2014.00028.
Henry, R. P. 2001. "Environmentally Mediated Carbonic Anhydrase Induction in the Gills of Euryhaline Crustaceans." *The Journal of Experimental Biology* 204 (Pt 5): 991–1002. https://doi.org/10.1242/jeb.204.5.991.
Hill, Jason, Erik D. Enbody, Mats E. Pettersson, C. Grace Sprehn, Dorte Bekkevold, Arild Folkvord, Linda Laikre, Gunnar Kleinau, Patrick Scheerer, and Leif Andersson. 2019. "Recurrent Convergent Evolution at Amino Acid Residue 261 in Fish Rhodopsin." *Proceedings of the National Academy of Sciences* 116 (37): 18473–78. https://doi.org/10.1073/pnas.1908332116.
Holtmann, Gudrun, Evert P. Bakker, Nobuyuki Uozumi, and Erhard Bremer. 2003. "KtrAB and KtrCD: Two K+ Uptake Systems in Bacillus Subtilis and Their Role in Adaptation to Hypertonicity." *Journal of Bacteriology*, February. https://doi.org/10.1128/JB.185.4.1289-1298.2003.
Hopkinson, Brian M., and Katherine A. Barbeau. 2012. "Iron Transporters in Marine Prokaryotic Genomes and Metagenomes." *Environmental Microbiology* 14 (1): 114–28. https://doi.org/10.1111/j.1462-2920.2011.02539.x.
Hsu, Jye-Lin, Hsuan-Cheng Chen, Hwei-Ling Peng, and Hwan-You Chang. 2008. "Characterization of the Histidine-Containing Phosphotransfer Protein B-Mediated Multistep Phosphorelay System in Pseudomonas Aeruginosa PAO1*." *Journal of Biological Chemistry* 283 (15): 9933–44. https://doi.org/10.1074/jbc.M708836200.
Hubloher, Josephine Joy, Sabine Zeidler, Pedro Lamosa, Helena Santos, Beate Averhoff, and Volker Müller. 2020. "Trehalose-6-Phosphate-Mediated Phenotypic Change in Acinetobacter Baumannii." *Environmental Microbiology* 22 (12): 5156–66. https://doi.org/10.1111/1462-2920.15148.
Iyer, Lakshminarayan M, Detlef D Leipe, Eugene V Koonin, and L Aravind. 2004. "Evolutionary History and Higher Order Classification of AAA+ ATPases." *Journal of Structural Biology*, AAA+ Proteins:  Based on Contributions to the 5th

International Conference on AAA+ Proteins  at Airlie House, Virginia, 146 (1): 11–31. https://doi.org/10.1016/j.jsb.2003.10.010.
Kamimura, Naofumi, and Eiji Masai. 2014. “The Protocatechuate 4,5-Cleavage Pathway: Overview and New Findings.” In *Biodegradative Bacteria: How Bacteria Degrade, Survive, Adapt, and Evolve*, edited by Hideaki Nojiri, Masataka Tsuda, Masao Fukuda, and Yoichi Kamagata, 207–26. Tokyo: Springer Japan. https://doi.org/10.1007/978-4-431-54520-0_10.
Kammler, M, C Schön, and K Hantke. 1993. “Characterization of the Ferrous Iron Uptake System of Escherichia Coli.” *Journal of Bacteriology* 175 (19): 6212–19. https://doi.org/10.1128/jb.175.19.6212-6219.1993.
Katsiou, E., C. M. Nickel, A. F. Garcia, and M. H. Tadros. 1999. “Molecular Analysis and Identification of the RadC Gene from the Phototrophic Bacterium Rhodobacter Capsulatus B10.” *Microbiological Research* 154 (3): 233–39. https://doi.org/10.1016/S0944-5013(99)80020-2.
Kędzierska, Barbara, Lu-Yun Lian, and Finbarr Hayes. 2007. “Toxin–Antitoxin Regulation: Bimodal Interaction of YefM–YoeB with Paired DNA Palindromes Exerts Transcriptional Autorepression.” *Nucleic Acids Research* 35 (1): 325–39. https://doi.org/10.1093/nar/gkl1028.
Kierzek, Elzbieta, and Ryszard Kierzek. 2003. “The Thermodynamic Stability of RNA Duplexes and Hairpins Containing N6-alkyladenosines and 2-methylthio-N6-alkyladenosines.” *Nucleic Acids Research* 31 (15): 4472–80. https://doi.org/10.1093/nar/gkg633.
Kim, Hyojoong, Seunghwan Lee, and Yikweon Jang. 2011. “Macroevolutionary Patterns in the Aphidini Aphids (Hemiptera: Aphididae): Diversification, Host Association, and Biogeographic Origins.” *PLOS ONE* 6 (9): e24749. https://doi.org/10.1371/journal.pone.0024749.
Kino, Kuniki, Toshinobu Arai, and Yasuhiro Arimura. 2011. “Poly-α-Glutamic Acid Synthesis Using a Novel Catalytic Activity of RimK from Escherichia Coli K-12.” *Applied and Environmental Microbiology* 77 (6): 2019–25. https://doi.org/10.1128/AEM.02043-10.
Kotakis, Christos, Eleftheria Theodoropoulou, Konstantinos Tassis, Charalambos Oustamanolakis, Nikolaos E. Ioannidis, and Kiriakos Kotzabasis. 2014. “Putrescine, a Fast-Acting Switch for Tolerance against Osmotic Stress.” *Journal of Plant Physiology* 171 (2): 48–51. https://doi.org/10.1016/j.jplph.2013.09.015.
Kuo, Chih-Horng, and Howard Ochman. 2009. “Inferring Clocks When Lacking Rocks: The Variable Rates of Molecular Evolution in Bacteria.” *Biology Direct* 4 (September): 35. https://doi.org/10.1186/1745-6150-4-35.
Lanclos, V. Celeste, Anna N. Rasmussen, Conner Y. Kojima, Chuankai Cheng, Michael W. Henson, Brant C. Faircloth, Christopher A. Francis, and J. Cameron Thrash. 2022. “Ecophysiology and Genomics of the Brackish Water Adapted SAR11 Subclade IIIa.” bioRxiv. https://doi.org/10.1101/2022.08.02.502558.
Levina, N., S. Tötemeyer, N. R. Stokes, P. Louis, M. A. Jones, and I. R. Booth. 1999. “Protection of Escherichia Coli Cells against Extreme Turgor by Activation of MscS and MscL Mechanosensitive Channels: Identification of Genes Required for MscS Activity.” *The EMBO Journal* 18 (7): 1730. https://doi.org/10.1093/emboj/18.7.1730.
Levine, Timothy P. 2019. “Remote Homology Searches Identify Bacterial Homologues of Eukaryotic Lipid Transfer Proteins, Including Chorein-N Domains in TamB and AsmA and Mdm31p.” *BMC Molecular and Cell Biology* 20 (1): 43. https://doi.org/10.1186/s12860-019-0226-z.

Li, Hong, Huimin Xu, David E. Graham, and Robert H. White. 2003. "Glutathione Synthetase Homologs Encode α-l-Glutamate Ligases for Methanogenic Coenzyme F420 and Tetrahydrosarcinapterin Biosyntheses." *Proceedings of the National Academy of Sciences of the United States of America* 100 (17): 9785–90. https://doi.org/10.1073/pnas.1733391100.

Lipscomb, John D. 2008. "Mechanism of Extradiol Aromatic Ring-Cleaving Dioxygenases." *Current Opinion in Structural Biology*, Catalysis and regulation / Proteins, 18 (6): 644–49. https://doi.org/10.1016/j.sbi.2008.11.001.

Little, Richard H., Lucia Grenga, Gerhard Saalbach, Alexandra M. Howat, Sebastian Pfeilmeier, Eleftheria Trampari, and Jacob G. Malone. 2016. "Adaptive Remodeling of the Bacterial Proteome by Specific Ribosomal Modification Regulates Pseudomonas Infection and Niche Colonisation." *PLoS Genetics* 12 (2): e1005837. https://doi.org/10.1371/journal.pgen.1005837.

Lo, N. 2003. "Evidence for Cocladogenesis Between Diverse Dictyopteran Lineages and Their Intracellular Endosymbionts." *Molecular Biology and Evolution* 20 (6): 907–13. https://doi.org/10.1093/molbev/msg097.

Mageeney, Catherine M, Britney Y Lau, Julian M Wagner, Corey M Hudson, Joseph S Schoeniger, Raga Krishnakumar, and Kelly P Williams. 2020. "New Candidates for Regulated Gene Integrity Revealed through Precise Mapping of Integrative Genetic Elements." *Nucleic Acids Research* 48 (8): 4052–65. https://doi.org/10.1093/nar/gkaa156.

Maguire, Michael E. 2006. "Magnesium Transporters: Properties, Regulation and Structure." *Frontiers in Bioscience: A Journal and Virtual Library* 11 (September): 3149–63. https://doi.org/10.2741/2039.

Marlovits, Thomas C., Winfried Haase, Christian Herrmann, Stephen G. Aller, and Vinzenz M. Unger. 2002. "The Membrane Protein FeoB Contains an Intramolecular G Protein Essential for Fe(II) Uptake in Bacteria." *Proceedings of the National Academy of Sciences* 99 (25): 16243–48. https://doi.org/10.1073/pnas.242338299.

Martorana, Alessandra M., Paola Sperandeo, Alessandra Polissi, and Gianni Dehò. 2011. "Complex Transcriptional Organization Regulates an Escherichia Coli Locus Implicated in Lipopolysaccharide Biogenesis." *Research in Microbiology* 162 (5): 470–82. https://doi.org/10.1016/j.resmic.2011.03.007.

Melville, Stephen, and Lisa Craig. 2013. "Type IV Pili in Gram-Positive Bacteria." *Microbiology and Molecular Biology Reviews* 77 (3): 323–41. https://doi.org/10.1128/MMBR.00063-12.

Merlin, Christophe, Millicent Masters, Sean McAteer, and Andrew Coulson. 2003. "Why Is Carbonic Anhydrase Essential to Escherichia Coli?" *Journal of Bacteriology* 185 (21): 6415–24. https://doi.org/10.1128/JB.185.21.6415-6424.2003.

Metzger, Lisa C., Noémie Matthey, Candice Stoudmann, Esther J. Collas, and Melanie Blokesch. 2019. "Ecological Implications of Gene Regulation by TfoX and TfoY among Diverse Vibrio Species." *Environmental Microbiology* 21 (7): 2231–47. https://doi.org/10.1111/1462-2920.14562.

Miller-Fleming, Leonor, Viridiana Olin-Sandoval, Kate Campbell, and Markus Ralser. 2015. "Remaining Mysteries of Molecular Biology: The Role of Polyamines in the Cell." *Journal of Molecular Biology* 427 (21): 3389–3406. https://doi.org/10.1016/j.jmb.2015.06.020.

Misra, R., and Y. Miao. 1995. "Molecular Analysis of AsmA, a Locus Identified as the Suppressor of OmpF Assembly Mutants of Escherichia Coli K-12." *Molecular Microbiology* 16 (4): 779–88. https://doi.org/10.1111/j.1365-2958.1995.tb02439.x.

Moore, R. C., and S. M. Boyle. 1991. "Cyclic AMP Inhibits and Putrescine Represses Expression of the SpeA Gene Encoding Biosynthetic Arginine Decarboxylase in Escherichia Coli." *Journal of Bacteriology* 173 (12): 3615–21. https://doi.org/10.1128/jb.173.12.3615-3621.1991.

Moran, Nancy A., Mark A. Munson, Paul Baumann, and Hajime Ishikawa. 1993. "A Molecular Clock in Endosymbiotic Bacteria Is Calibrated Using the Insect Hosts." *Proceedings of the Royal Society of London. Series B: Biological Sciences* 253 (1337): 167–71. https://doi.org/10.1098/rspb.1993.0098.

Moran, Nancy A., Phat Tran, and Nicole M. Gerardo. 2005. "Symbiosis and Insect Diversification: An Ancient Symbiont of Sap-Feeding Insects from the Bacterial Phylum Bacteroidetes." *Applied and Environmental Microbiology* 71 (12): 8802–10. https://doi.org/10.1128/AEM.71.12.8802-8810.2005.

Mueller, Lukas A., Ursula Hinz, and Jean-Pierre Zryd. 1997. "The Formation of Betalamic Acid and Muscaflavin by Recombinant Dopa-Dioxygenase from Amanita." *Phytochemistry* 44 (4): 567–69. https://doi.org/10.1016/S0031-9422(96)00625-5.

Muñoz-García, Andrea, Orson Mestanza, Juan Pablo Isaza, Ingrid Figueroa-Galvis, and Javier Vanegas. 2019. "Influence of Salinity on the Degradation of Xenobiotic Compounds in Rhizospheric Mangrove Soil." *Environmental Pollution* 249 (June): 750–57. https://doi.org/10.1016/j.envpol.2019.03.056.

Namugenyi, Sarah B., Alisha M. Aagesen, Sarah R. Elliott, and Anna D. Tischler. 2017. "Mycobacterium Tuberculosis PhoY Proteins Promote Persister Formation by Mediating Pst/SenX3-RegX3 Phosphate Sensing." *MBio* 8 (4): e00494-17. https://doi.org/10.1128/mBio.00494-17.

Nanatani, Kei, Toshiaki Shijuku, Yousuke Takano, Lalu Zulkifli, Tomoko Yamazaki, Akira Tominaga, Satoshi Souma, et al. 2015. "Comparative Analysis of Kdp and Ktr Mutants Reveals Distinct Roles of the Potassium Transporters in the Model Cyanobacterium Synechocystis Sp. Strain PCC 6803." *Journal of Bacteriology* 197 (4): 676–87. https://doi.org/10.1128/JB.02276-14.

Noda, Y, S Nishikawa, K Shiozuka, H Kadokura, H Nakajima, K Yoda, Y Katayama, N Morohoshi, T Haraguchi, and M Yamasaki. 1990. "Molecular Cloning of the Protocatechuate 4,5-Dioxygenase Genes of Pseudomonas Paucimobilis." *Journal of Bacteriology* 172 (5): 2704–9. https://doi.org/10.1128/jb.172.5.2704-2709.1990.

Payandeh, Jian, Roland Pfoh, and Emil F. Pai. 2013. "The Structure and Regulation of Magnesium Selective Ion Channels." *Biochimica Et Biophysica Acta* 1828 (11): 2778–92. https://doi.org/10.1016/j.bbamem.2013.08.002.

Price, G. Dean, Fiona J. Woodger, Murray R. Badger, Susan M. Howitt, and Loraine Tucker. 2004. "Identification of a SulP-Type Bicarbonate Transporter in Marine Cyanobacteria." *Proceedings of the National Academy of Sciences* 101 (52): 18228–33. https://doi.org/10.1073/pnas.0405211101.

Prieto, Ana I., Sara B. Hernández, Ignacio Cota, M. Graciela Pucciarelli, Yuri Orlov, Francisco Ramos-Morales, Francisco García-del Portillo, and Josep Casadesús. 2009. "Roles of the Outer Membrane Protein AsmA of Salmonella Enterica in the Control of MarRAB Expression and Invasion of Epithelial Cells." *Journal of Bacteriology* 191 (11): 3615–22. https://doi.org/10.1128/JB.01592-08.

Qi, Y., Y. Kobayashi, and F. M. Hulett. 1997. "The Pst Operon of Bacillus Subtilis Has a Phosphate-Regulated Promoter and Is Involved in Phosphate Transport but Not in Regulation of the Pho Regulon." *Journal of Bacteriology* 179 (8): 2534–39. https://doi.org/10.1128/jb.179.8.2534-2539.1997.

Ramachandran, Arthi, Susan McLatchie, and David A. Walsh. 2021. "A Novel Freshwater to Marine Evolutionary Transition Revealed within Methylophilaceae Bacteria from the Arctic Ocean." *MBio* 12 (3): e01306-21. https://doi.org/10.1128/mBio.01306-21.

Rodriguez-Navarro, A., M. R. Blatt, and C. L. Slayman. 1986. "A Potassium-Proton Symport in Neurospora Crassa." *The Journal of General Physiology* 87 (5): 649–74. https://doi.org/10.1085/jgp.87.5.649.

Ruhal, Rohit, Rashmi Kataria, and Bijan Choudhury. 2013. "Trends in Bacterial Trehalose Metabolism and Significant Nodes of Metabolic Pathway in the Direction of Trehalose Accumulation." *Microbial Biotechnology* 6 (5): 493–502. https://doi.org/10.1111/1751-7915.12029.

Salzer, Ralf, Timo Kern, Friederike Joos, and Beate Averhoff. 2016. "The Thermus Thermophilus ComEA/ComEC Operon Is Associated with DNA Binding and Regulation of the DNA Translocator and Type IV Pili." *Environmental Microbiology* 18 (1): 65–74. https://doi.org/10.1111/1462-2920.12820.

Seitz, Patrick, and Melanie Blokesch. 2013. "Cues and Regulatory Pathways Involved in Natural Competence and Transformation in Pathogenic and Environmental Gram-Negative Bacteria." *FEMS Microbiology Reviews* 37 (3): 336–63. https://doi.org/10.1111/j.1574-6976.2012.00353.x.

Seitz, Patrick, Hassan Pezeshgi Modarres, Sandrine Borgeaud, Roman D. Bulushev, Lorenz J. Steinbock, Aleksandra Radenovic, Matteo Dal Peraro, and Melanie Blokesch. 2014. "ComEA Is Essential for the Transfer of External DNA into the Periplasm in Naturally Transformable Vibrio Cholerae Cells." *PLoS Genetics* 10 (1): e1004066. https://doi.org/10.1371/journal.pgen.1004066.

Sergiev, Petr V., Anna Ya. Golovina, Ilya A. Osterman, Michail V. Nesterchuk, Olga V. Sergeeva, Anastasia A. Chugunova, Sergey A. Evfratov, et al. 2016. "N6-Methylated Adenosine in RNA: From Bacteria to Humans." *Journal of Molecular Biology*, Ribosomes Structure and Mechanisms in Regulation of Protein Synthesis (I), 428 (10, Part B): 2134–45. https://doi.org/10.1016/j.jmb.2015.12.013.

Sestok, Alex E., Sean M. O'Sullivan, and Aaron T. Smith. 2022. "A General Protocol for the Expression and Purification of the Intact Transmembrane Transporter FeoB." *Biochimica Et Biophysica Acta. Biomembranes* 1864 (9): 183973. https://doi.org/10.1016/j.bbamem.2022.183973.

Shan, Shan, Haowei Min, Ting Liu, Dunquan Jiang, and Zihe Rao. 2016. "Structural Insight into Dephosphorylation by Trehalose 6-Phosphate Phosphatase (OtsB2) from Mycobacterium Tuberculosis." *The FASEB Journal* 30 (12): 3989–96. https://doi.org/10.1096/fj.201600463R.

Shcherbakov, Dmitry. 2002. "The 270 Million Year History of Auchenorrhyncha (Homoptera)." *Denisia* 4 (January): 29–36.

Silale, Augustinas, Susan M. Lea, and Ben C. Berks. 2021. "The DNA Transporter ComEC Has Metal-Dependent Nuclease Activity That Is Important for Natural Transformation." *Molecular Microbiology* 116 (2): 416–26. https://doi.org/10.1111/mmi.14720.

Smith, K. S., and J. G. Ferry. 2000. "Prokaryotic Carbonic Anhydrases." *FEMS Microbiology Reviews* 24 (4): 335–66. https://doi.org/10.1111/j.1574-6976.2000.tb00546.x.

Springael, Dirk, and Eva M Top. 2004. "Horizontal Gene Transfer and Microbial Adaptation to Xenobiotics: New Types of Mobile Genetic Elements and Lessons from Ecological Studies." *Trends in Microbiology* 12 (2): 53–58. https://doi.org/10.1016/j.tim.2003.12.010.

Stetsenko, Artem, and Albert Guskov. 2020. "Cation Permeability in CorA Family of Proteins." *Scientific Reports* 10 (1): 840. https://doi.org/10.1038/s41598-020-57869-z.

Stevenson, Begoña, Elizabeth E. Wyckoff, and Shelley M. Payne. 2016. "Vibrio Cholerae FeoA, FeoB, and FeoC Interact To Form a Complex." *Journal of Bacteriology* 198 (7): 1160–70. https://doi.org/10.1128/JB.00930-15.

Strøm, A. R., and I. Kaasen. 1993. "Trehalose Metabolism in Escherichia Coli: Stress Protection and Stress Regulation of Gene Expression." *Molecular Microbiology* 8 (2): 205–10. https://doi.org/10.1111/j.1365-2958.1993.tb01564.x.

Suzuki, Shingo, Ali Ferjani, Iwane Suzuki, and Norio Murata. 2004. "The SphS-SphR Two Component System Is the Exclusive Sensor for the Induction of Gene Expression in Response to Phosphate Limitation in Synechocystis." *The Journal of Biological Chemistry* 279 (13): 13234–40. https://doi.org/10.1074/jbc.M313358200.

Szklarczyk, Damian, Annika L. Gable, David Lyon, Alexander Junge, Stefan Wyder, Jaime Huerta-Cepas, Milan Simonovic, et al. 2019. "STRING V11: Protein-Protein Association Networks with Increased Coverage, Supporting Functional Discovery in Genome-Wide Experimental Datasets." *Nucleic Acids Research* 47 (D1): D607–13. https://doi.org/10.1093/nar/gky1131.

Szumanski, M B, and S M Boyle. 1992. "Influence of Cyclic AMP, Agmatine, and a Novel Protein Encoded by a Flanking Gene on SpeB (Agmatine Ureohydrolase) in Escherichia Coli." *Journal of Bacteriology* 174 (3): 758–64. https://doi.org/10.1128/jb.174.3.758-764.1992.

Tao, T., M. D. Snavely, S. G. Farr, and M. E. Maguire. 1995. "Magnesium Transport in Salmonella Typhimurium: MgtA Encodes a P-Type ATPase and Is Regulated by Mg2+ in a Manner Similar to That of the MgtB P-Type ATPase." *Journal of Bacteriology*, May. https://doi.org/10.1128/jb.177.10.2654-2662.1995.

Tascón, Igor, Joana S. Sousa, Robin A. Corey, Deryck J. Mills, David Griwatz, Nadine Aumüller, Vedrana Mikusevic, Phillip J. Stansfeld, Janet Vonck, and Inga Hänelt. 2020. "Structural Basis of Proton-Coupled Potassium Transport in the KUP Family." *Nature Communications* 11 (1): 626. https://doi.org/10.1038/s41467-020-14441-7.

Thao, M. L., N. A. Moran, P. Abbot, E. B. Brennan, D. H. Burckhardt, and P. Baumann. 2000. "Cospeciation of Psyllids and Their Primary Prokaryotic Endosymbionts." *Applied and Environmental Microbiology* 66 (7): 2898–2905. https://doi.org/10.1128/AEM.66.7.2898-2905.2000.

Trchounian, A., and H. Kobayashi. 1999. "Kup Is the Major K+ Uptake System in Escherichia Coli upon Hyper-Osmotic Stress at a Low PH." *FEBS Letters* 447 (2–3): 144–48. https://doi.org/10.1016/s0014-5793(99)00288-4.

Tsagogiannis, Epameinondas, Elpiniki Vandera, Alexandra Primikyri, Stamatia Asimakoula, Andreas G. Tzakos, Ioannis P. Gerothanassis, and Anna-Irini Koukkou. 2021. "Characterization of Protocatechuate 4,5-Dioxygenase from Pseudarthrobacter Phenanthrenivorans Sphe3 and In Situ Reaction Monitoring in the NMR Tube." *International Journal of Molecular Sciences* 22 (17): 9647. https://doi.org/10.3390/ijms22179647.

Västermark, Åke, and Milton H. Saier. 2014. "The Involvement of Transport Proteins in Transcriptional and Metabolic Regulation." *Current Opinion in Microbiology* 0 (April): 8–15. https://doi.org/10.1016/j.mib.2014.01.002.

Vigil-Stenman, Theoden, Karolina Ininbergs, Birgitta Bergman, and Martin Ekman. 2017. "High Abundance and Expression of Transposases in Bacteria from the Baltic Sea." *The ISME Journal* 11 (11): 2611–23. https://doi.org/10.1038/ismej.2017.114.

Walsh, David A, Josine Lafontaine, and Hans-Peter Grossart. 2013. "On the Eco-Evolutionary Relationships of Fresh and Salt Water Bacteria and the Role of Gene Transfer in Their Adaptation." In *Lateral Gene Transfer in Evolution*, edited by Uri

Gophna, 55–77. New York, NY: Springer. https://doi.org/10.1007/978-1-4614-7780-8_3.
Webb, D. C., H. Rosenberg, and G. B. Cox. 1992. “Mutational Analysis of the Escherichia Coli Phosphate-Specific Transport System, a Member of the Traffic ATPase (or ABC) Family of Membrane Transporters. A Role for Proline Residues in Transmembrane Helices.” *The Journal of Biological Chemistry* 267 (34): 24661–68.
Wen, Yi, Jing Feng, David R. Scott, Elizabeth A Marcus, and George Sachs. 2009. “The PH-Responsive Regulon of HP0244 (FlgS), the Cytoplasmic Histidine Kinase of Helicobacter Pylori.” *Journal of Bacteriology* 191 (2): 449–60. https://doi.org/10.1128/JB.01219-08.
Witt, G. 1995. “Polycyclic Aromatic Hydrocarbons in Water and Sediment of the Baltic Sea.” *Marine Pollution Bulletin*, Selected Papers from the International Conference on Marine Pollution and Ecotoxicology, 31 (4): 237–48. https://doi.org/10.1016/0025-326X(95)00174-L.
Wösten, Marc M. S. M., Jaap A. Wagenaar, and Jos P. M. van Putten. 2004. “The FlgS/FlgR Two-Component Signal Transduction System Regulates the Fla Regulon in Campylobacter Jejuni.” *The Journal of Biological Chemistry* 279 (16): 16214–22. https://doi.org/10.1074/jbc.M400357200.
Yu, Jie, Bing Zhang, Yixiao Zhang, Cong-Qiao Xu, Wei Zhuo, Jingpeng Ge, Jun Li, Ning Gao, Yang Li, and Maojun Yang. 2018. “A Binding-Block Ion Selective Mechanism Revealed by a Na/K Selective Channel.” *Protein & Cell* 9 (7): 629–39. https://doi.org/10.1007/s13238-017-0465-8.
Zamakhaev, Mikhail, Ivan Tsyganov, Larisa Nesterova, Anna Akhova, Artem Grigorov, Julia Bespyatykh, Tatyana Azhikina, Alexander Tkachenko, and Mikhail Shumkov. 2020. “Mycolicibacterium Smegmatis Possesses Operational Agmatinase but Contains No Detectable Polyamines.” *International Journal of Mycobacteriology* 9 (2): 138–43. https://doi.org/10.4103/ijmy.ijmy_48_20.
Zeidler, Sabine, Josephine Hubloher, Kim Schabacker, Pedro Lamosa, Helena Santos, and Volker Müller. 2017. “Trehalose, a Temperature- and Salt-Induced Solute with Implications in Pathobiology of Acinetobacter Baumannii.” *Environmental Microbiology* 19 (12): 5088–99. https://doi.org/10.1111/1462-2920.13987.
Zhang, Hao, Susumu Yoshizawa, Ying Sun, Yongjie Huang, Xiao Chu, José M. González, Jarone Pinhassi, and Haiwei Luo. 2019. “Repeated Evolutionary Transitions of Flavobacteria from Marine to Non-Marine Habitats.” *Environmental Microbiology* 21 (2): 648–66. https://doi.org/10.1111/1462-2920.14509.
Zolotarev, Alexander S., Meera Unnikrishnan, Boris E. Shmukler, Jeffrey S. Clark, David H. Vandorpe, Nikolaus Grigorieff, Eric J. Rubin, and Seth L. Alper. 2008. “Increased Sulfate Uptake by E. Coli Overexpressing the SLC26-Related SulP Protein Rv1739c from Mycobacterium Tuberculosis.” *Comparative Biochemistry and Physiology. Part A, Molecular & Integrative Physiology* 149 (3): 255–66. https://doi.org/10.1016/j.cbpa.2007.12.005.
